# A Novel Family of RNA-Binding Proteins Regulate Polysaccharide Metabolism in *Bacteroides thetaiotaomicron*

**DOI:** 10.1101/2021.04.27.441718

**Authors:** Amanda N.D. Adams, Muhammad S. Azam, Zachary A. Costliow, Xiangqian Ma, Patrick H. Degnan, Carin K. Vanderpool

**Author notes:** Correspondence to: Patrick H. Degnan,; or Carin K. Vanderpool.

## Abstract

Human gut microbiome composition is constantly changing, and diet is a major driver of these changes. Gut microbial species that persist in mammalian hosts for long periods of time must possess mechanisms for sensing and adapting to nutrient shifts to avoid being outcompeted. Global regulatory mechanisms mediated by RNA-binding proteins (RBPs) that govern responses to nutrient shifts have been characterized in Proteobacteria and Firmicutes but remain undiscovered in the Bacteroidetes. Here we report the identification of RBPs that are broadly distributed across the Bacteroidetes, with many genomes encoding multiple copies. Genes encoding these RBPs are highly expressed in many *Bacteroides* species. A purified RBP, RbpB, from *Bacteroides thetaiotaomicron* binds to single-stranded RNA *in vitro* with an affinity similar to other characterized regulatory RBPs. *B. thetaiotaomicron* mutants lacking RBPs show dramatic shifts in expression of polysaccharide utilization and capsular polysaccharide loci, suggesting that these RBPs may act as global regulators of polysaccharide metabolism. A *B. thetaiotaomicron* Δ*rbpB* mutant shows a growth defect on dietary sugars belonging to the raffinose family of oligosaccharides (RFOs). The Δ*rbpB* mutant had reduced expression of *BT1871*, encoding a predicted RFO-degrading melibiase, compared to the wild-type strain. Mutation of *BT1871* confirmed that the enzyme it encodes is essential for growth on melibiose and promotes growth on the RFOs raffinose and stachyose. Our data reveal that RbpB is required for optimal expression of *BT1871* and other polysaccharide-related genes, suggesting that we have identified an important new family of global regulatory proteins in the Bacteroidetes.

**Importance:** The human colon houses hundreds of bacterial species, including many belonging to the genus *Bacteroides,* that aid in breaking down our food to keep us healthy. *Bacteroides* have many genes responsible for breaking down different dietary carbohydrates and complex regulatory mechanisms ensure that specific genes are only expressed when the right carbohydrates are available. In this study, we discovered that *Bacteroides* use a family of RNA-binding proteins as global regulators to coordinate expression of carbohydrate utilization genes. The ability to turn different carbohydrate utilization genes on and off in response to changing nutrient conditions is critical for *Bacteroides* to live successfully in the gut, and thus the new regulators we have identified may be important for life in the host.

## Introduction

The human gut microbiome is an important player in host health, with diet being one of the principal drivers of gut microbial composition and function (1–3). Dietary carbohydrates, including complex polysaccharides and oligosaccharides are not readily absorbed by the host and reach the distal gut where they are broken down and metabolized by a consortium of microbes with diverse enzymatic capabilities (4).

Members of the dominant bacterial phylum Bacteroidetes can readily switch between carbohydrate types as they become available due to dozens of substrate-specific polysaccharide utilization loci (PUL) that encode proteins responsible for sensing and catabolizing diverse polysaccharides (5–7). Characterized PULs are tightly regulated by several distinct families of transcriptional regulators so that they are only abundantly expressed when their substrates are available (5, 8–17). However, accumulating evidence suggests that post-transcriptional regulation also plays an important part in gut colonization and preferential use of carbohydrates through control of PULs (18–20).

Post-transcriptional regulation can be mediated by multiple regulators. In *Bacteroides* species, the roles of small RNA (sRNA) regulators (19, 21) and other RNA regulatory elements like riboswitches (22–24) in control of carbohydrate and vitamin metabolism are beginning to be recognized. In well-studied Firmicutes and Proteobacteria, post-transcriptional regulation of carbon metabolism and other systems often occurs through the actions of sRNAs and their helper RNA chaperones (25).

Three of the most well-studied RNA chaperones include Hfq, CsrA, and ProQ. Collectively, these three RNA-binding proteins (RBPs) regulate the bulk of the RNA regulatory interactome in organisms like *Escherichia coli* and *Salmonella enterica*, and each RBP has its own distinct RNA targets (26, 27). Hfq in particular functions as a global post-transcriptional regulator of gene expression (28, 29). It binds to both mRNAs and sRNAs, facilitating their interactions through short stretches of complementarity (30, 31). These interactions result in a variety of different regulatory outcomes, primarily resulting from changes in translation initiation or mRNA stability (28). In many organisms, mutation of *hfq* causes global changes in gene expression and pleiotropic phenotypes (32–34).

Though regulatory RNA chaperones have not been characterized in the Bacteroidetes, post-transcriptional regulation has been implicated in control of gene expression in *Bacteroides* species (21, 35), including regulation of PUL expression (19). In particular, the cis-antisense PUL-associated sRNA DonS in *B. fragilis* as well as several PUL-associated cis-antisense sRNAs in *B. thetaiotaomicron* (19, 21) have been implicated in modulation of PUL function through repression of carbohydrate transporter gene expression. A recent study identified dozens of sRNAs encoded throughout the genome of *B. thetaiotaomicron* with many being PUL-associated, suggesting that sRNA-mediated regulation of PUL function may be a common phenomenon in *Bacteroides* (21). Additionally, there are a growing number of examples of regulatory effects in *Bacteroides* mediated by sequences in mRNA untranslated regions (UTRs) (18, 36) and these may be mediated by as yet unidentified sRNAs or RNA chaperones. To better understand the scope of RNA-mediated regulatory mechanisms in the Bacteroidetes, we sought to identify and characterize RBPs that may act as regulatory RNA chaperones.

Here we report the identification of a family of genes commonly found in Bacteroidetes genomes, which encode RBPs with a single RNA Recognition Motif 1 (RRM-1) domain. These genes are conserved, often exist in multiple copies per genome, and are highly expressed in many human gut *Bacteroides* isolates. We demonstrate that a member of this family, RbpB, is a single-stranded (ss) RNA-binding protein that binds with some specificity and affinities similar to other characterized RNA chaperones. *B. thetaiotaomicron* mutants lacking one or more of these RBPs have large-scale changes to their transcriptomes compared to the wild-type strain, with genes belonging to PUL and capsular polysaccharide (CPS) loci being the most differentially regulated. *B. thetaiotaomicron rbpB* mutants have growth defects on the common dietary plant sugars raffinose family oligosaccharides due to decreased expression of *BT1871*, an essential melibiase encoded in PUL24. Our findings suggest that this family of RBPs play an important role in global regulation of polysaccharide metabolism in *Bacteroides*.

## Results

### Identification of a conserved family of RNA-binding proteins in the phylum Bacteroidetes

To identify putative RNA-binding proteins that may act as global regulators in the Bacteroidetes, we compared a set of 313 human gut-associated microbial genomes representing major phyla commonly found in gut microbial communities (37). We first searched for canonical RNA chaperones – Hfq, ProQ, and CsrA, which are involved in post-transcriptional regulation of gene expression in Proteobacteria. Using hidden Markov models (HMMs) with trusted cutoffs, we identified Hfq in 23% (72/313) of the genomes (a total of 79 Hfq homologs) mostly in the Proteobacteria, although there were some identified in Firmicutes genomes (Fig.1A and Dataset S1). ProQ homologs (a total of 40 across 36 genomes) were entirely restricted to the Proteobacteria, with the majority being found in γ-proteobacterial genomes. CsrA homologs were identified in 25% (79/313) of genomes with a total of 93 CsrA homologs distributed across the Proteobacteria and Firmicutes. We did not identify any Hfq, ProQ, or CsrA homologs among Bacteroidetes genomes suggesting that if RNA chaperone regulators are present in this phylum, they do not belong to these canonical families.

**Figure 1:**
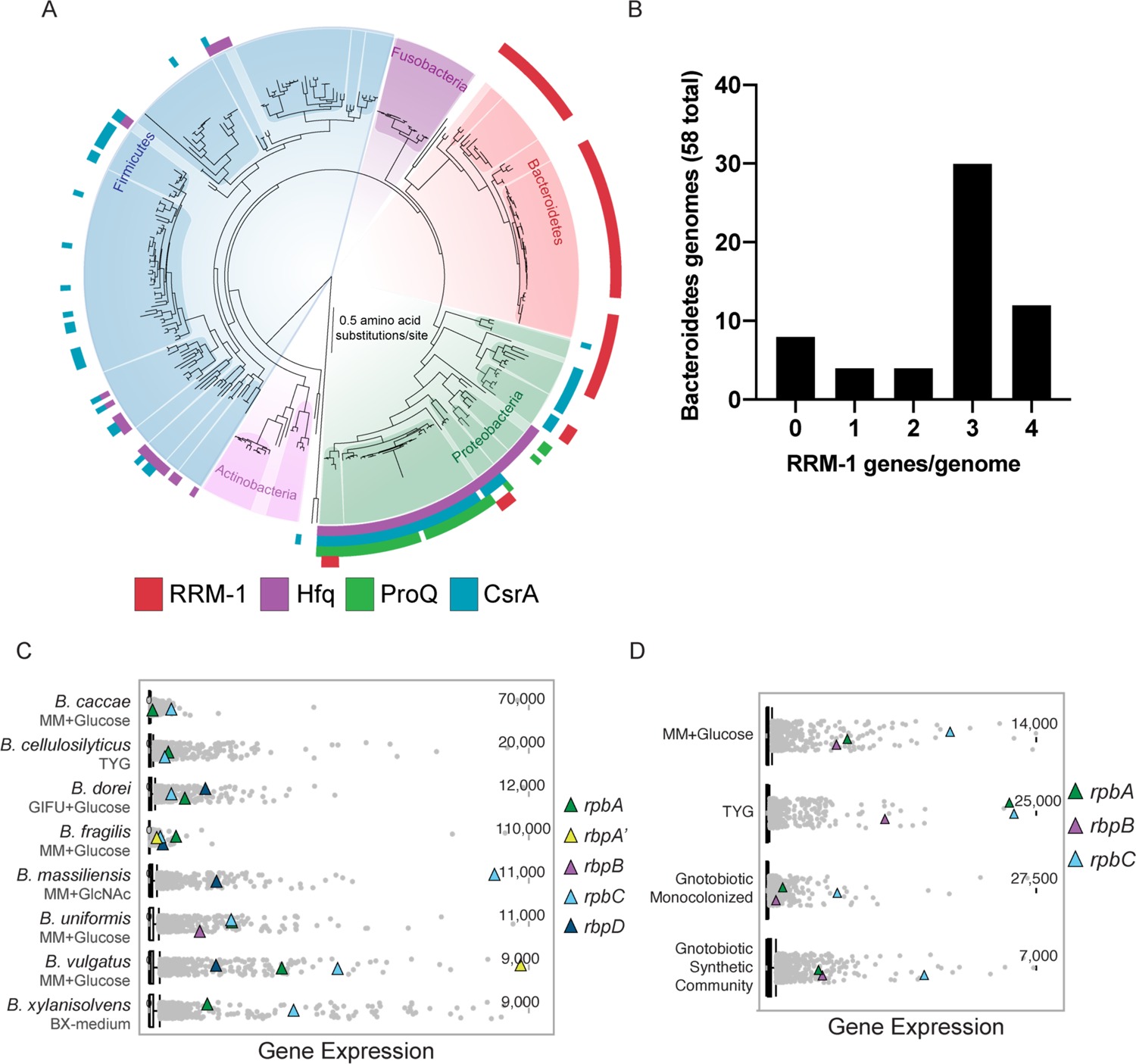
RRM-1 is a conserved, abundantly expressed RNA-binding domain in gut bacteria. (A) Maximum likelihood phylogenetic species tree of 313 human gut-associated microbial genomes. Colored bars indicate presence of at least one copy of the indicated RNA regulatory proteins in a given genome. (B) Histogram of total RRM-1 genes per genome in the 58 Bacteroidetes genomes represented in (A). (C-D) RNA-seq expression plots of all genes in publicly available transcriptomes for various *Bacteroides* species (C) or *B. thetaiotaomicron* only in various growth conditions (D). Grey dots represent a single gene and triangles *rbp* genes. The top 10% of expressed genes lie above the whiskers.

To identify other putative RNA chaperones in our model organism, we searched the *B. thetaiotaomicron* VPI-5482 genome for proteins with conserved RNA-binding domains. This yielded three intriguing candidates comprised of a single RNA Recognition Motif 1 (RRM-1; PF00076) domain, here named RbpA (BT0784), RbpB (BT1887), and RbpC (BT3840). The small, ∼70 amino acid RRM-1 domain is one of the most common RNA-binding domains in eukaryotes where it is typically found in multidomain proteins involved in post-transcriptional RNA processing events including regulation of RNA stability, translation, and turnover (38). Though poorly characterized, many bacterial genomes appear to encode RRM-1 domain-containing proteins (39, 40). Given the characterized roles of RRM-1 domain-containing proteins in post-transcriptional RNA regulatory processes, we chose to focus on these homologs for further characterization.

Expanding our search for RRM-1 domain-containing proteins to our larger set of gut microbial genomes identified homologs of *B. thetaiotaomicron* RBPs in 69 of 313 genomes (Fig. 1A and Dataset S1). These proteins were widely distributed among Bacteroidetes genomes accounting for 86% (149/174) of the total number of RRM-1 domain proteins identified. We also identified homologs in a small subset of Proteobacterial genomes (Fig. 1A, Dataset S1). In contrast to eukaryotes, the bacterial RRM-1 proteins we identified are small, single-domain proteins ranging in size from 60-132 amino acids (aa), with the majority being 80-100 aa. Each protein contains a single ferredoxin-like fold RRM-1 motif followed by a predicted disordered C-terminus of varying lengths. This structure is reminiscent of the disordered C-termini of Hfq and other RNA chaperones that plays a role in RNA-binding and cycling among various binding partners (41–45). CsrA, Hfq, and ProQ homologs were largely encoded in single copy with only a few instances of more than one copy in individual genomes (Dataset S1). In contrast, RRM-1 genes frequently occurred in multiple copies per genome in Bacteroidetes genomes (Table S1, Fig. 1B). Of the Bacteroidetes genomes we analyzed, 50 of 58 contained one to four copies of genes encoding RRM-1 domain proteins, with the majority of genomes containing three (Fig. 1B).

To analyze phylogenetic relationships among novel RRM-1 domain proteins found in Bacteroidetes genomes, we used MCL (Markov cluster algorithm) with a 70% amino acid identity cutoff and compared the resulting clusters to a species phylogeny of the Bacteroidetes (Dataset S1). Clustering was chosen because the extent of divergence among the homologs and their short lengths makes phylogenetic reconstruction unreliable. The clustering revealed a complicated history of divergence and duplication resulting in 10 clusters designated *rbpA–rbpJ.* It is notable that the three loci represented in *B. thetaiotaomicron*, *rbpA, rbpB,* and *rbpC* represent the most widespread clusters. Even within these clusters we identified evidence of likely duplications or horizontal gene transfer among particular lineages, resulting in genomes that encode two genes belonging to a single cluster. For example, the *Bacteroides fragilis* 3_1_2 genome contains *rbpA* and *rbpA’* which share 84% amino acid identity.

The well-characterized RNA chaperone Hfq is an abundant protein in Proteobacteria (46). To determine whether genes encoding *Bacteroides* RBPs show similarly high levels of expression, we analyzed available RNA-seq data for *B. thetaiotaomicron* (generated by us (see Materials and Methods) and others (23)) and eight additional species – *Bacteroides caccae* (47)*, Bacteroides cellulosyliticus* (48)*, Bacteroides dorei* (49)*, Bacteroides massilensis* (18)*, Bacteroides uniformis* (24)*, Bacteroides vulgatus* (24), and *Bacteroides xylanisolvens* (50) that were grown under a variety of *in vitro* conditions. Virtually all of the genes encoding RBP homologs were highly expressed in these datasets. Most *Bacteroides rbp* genes (represented by colored triangles, Fig. 1C, 1D) were expressed at levels placing them among the top 10% of most highly expressed genes (represented by grey dots, Fig. 1C, 1D). Looking specifically at *B. thetaiotaomicron rbpA, rbpB,* and *rbpC*, we observed that these genes are highly expressed both *in vitro* (minimal medium with glucose and TYG medium) and *in vivo* in monocolonized mice (51) or mice colonized with a synthetic consortium (47) (Fig. 1D).

### *B. thetaiotaomicron* RbpB is a single-stranded RNA-binding protein

To test the RNA-binding activity of a representative of this family of *Bacteroides* RBPs, we conducted electrophoretic mobility shift assays (EMSAs). We overexpressed and purified *B. thetaiotaomicron* RbpB and tested binding to a series of *in vitro* transcribed single-stranded RNA (ssRNA) “pentaprobes” (52). The twelve ssRNA pentaprobes are each 100-nucleotides (nts) in length and collectively contain all possible 5-nt sequence combinations (Table S1). RbpB shifted ten of the twelve probes to varying degrees, indicating that RbpB binds ssRNA *in vitro* and suggesting that it does so with some degree of sequence specificity (Figs. 2A and S1A). RbpB showed no evidence of binding to two of the pentaprobes PP6 and PP12 (Figs. S1A and 2B).

**Figure 2:**
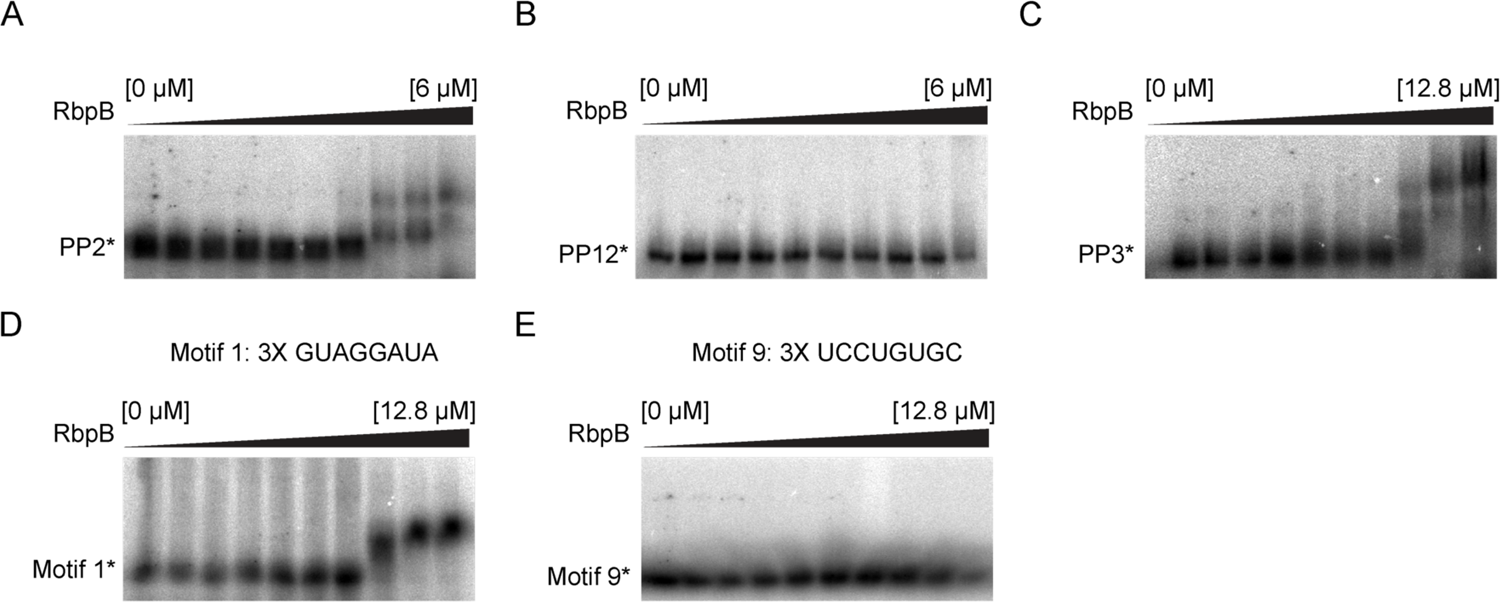
RbpB is a ssRNA-binding protein. (A-B) RbpB-pentaprobe EMSAs for pentaprobes 2 (A) and 12 (B) (PP2 and PP12, respectively). RbpB [μM] increases from left to right as follows: 0, 0.02, 0.05, 0.09, 0.19, 0.38, 0.75, 1.50, 3.00, 6.00. (C) Pentaprobe 3 repeat EMSA with RbpB [μM] increasing from left to right in the gel as follows: 0, 0.05, 0.10, 0.20, 0.40, 0.80, 1.60, 3.20, 6.40, 12.80. (D-E) EMSAs of a 3x repeat of MEME motif 1 (D) or motif 9 (E) with RbpB increasing from left to right as in (C). Asterisk * indicates the unbound radiolabeled pentaprobe.

Probes PP2 and PP3 shifted at lower concentrations of RbpB compared with other probes (Figs. 2A and S1A). The calculated *K_D_* of RbpB binding to PP3 is 10.5 µM (Fig. 2C), a dissociation constant similar to those previously reported for RRM domains (38, 53). RRM domains can interact with a variable number of nts in the binding pocket, with binding motifs that are typically 5-8 nts in length (54). To identify candidate RbpB binding motifs in the pentaprobes, we used MEME motif discovery tool (55) to identify sequence motifs (<9-nt in length) that occurred in RbpB-binding pentaprobes but were absent in non-binding pentaprobes (Fig. S1B). MEME identified 10 such motifs in RbpB-binding pentaprobe sequences. Motif 1, comprised of G, U, and A residues was the most common motif found exclusively in the bound pentaprobes (Figs. 2D and S1B). A C/U-rich motif, motif 9 (Fig. 2E and S1B), was present in probes PP2 and PP3, which bound RbpB with higher affinity than other pentaprobes (Fig. 2A and C and S1A). To test whether RbpB would bind specifically to motif 1 (5’-GUAGGAUA-3’) or motif 9 (5’UCCUGUGC-3’), we conducted EMSAs using new RNA oligonucleotide probes containing three repeats of each motif. RbpB shifted the probe containing three copies of motif 1 with a *K_D_* of 5.1 µM (Fig. 2D). In contrast, RbpB did not shift a probe containing three copies of motif 9 (Fig. 2E). Overall, these results demonstrate that RbpB binds ssRNA with some degree of specificity at affinities comparable to known RNA-binding proteins (56–58).

### Loss of RBPs leads to altered expression of PUL and CPS loci

To assess possible functions of RBPs in *B. thetaiotaomicron*, we made mutant strains lacking *rbpA* and *rbpB*. We generated three strains: Δ*rbpA (BT0784)*, Δ*rbpB (BT1887),* and Δ*rbpAΔrbpB.* We were unable to generate a Δ*rbpC* mutant. We performed RNA-seq on RNA samples from Δ*rbpA,* Δ*rbpB,* and Δ*rbpAΔrbpB* strains grown to mid-log or stationary phase in rich medium (TYG). Among protein coding genes, 12.3% (587/4,778) were significantly differentially regulated (q-value <0.06, log_2_ fold-change of ≥+1 or ≤-1) in at least one condition (Dataset S2A), with Δ*rbpA*Δ*rbpB* having the greatest number of differentially regulated genes among the three mutants. To identify functional classes of differentially expressed genes, Gene Set Enrichment Analysis (GSEA) (59) was used with *B. thetaiotaomicron-*specific custom gene sets (see Materials and Methods). Differentially regulated genes that were not categorized in GSEA were further grouped according to Gene Ontology (Dataset S2A and Materials and Methods). Considering all differentially regulated genes across all strains, enriched functional groups included CPS loci, PULs, hypothetical proteins, transmembrane transport, redox activities, B-vitamin metabolism, transcription, translation, and a variety of other metabolic pathways (Fig. 3A). The largest functional group of differentially regulated genes was CPS genes, accounting for 17% (98/587) of differentially regulated genes across all six wild-type to mutant comparisons (Fig. 3A, B). CPS loci encode functions that produce the polysaccharide coats that surround the *Bacteroides* cell surface (10, 17, 60). Of the eight CPS loci in *B. thetaiotaomicron* VPI-5482, five were differentially regulated across the three mutant strains including CPS2, CPS4, CPS5, CPS6, and CPS7 loci (Fig. 3B). CPS4 and 6 loci were downregulated in all three mutants compared to wild-type in both conditions, whereas CPS2, 5, and 7 loci were upregulated in some mutants compared to wild-type in a subset of conditions. CPS2 was upregulated in both Δ*rbpA* and Δ*rbpB* mutants in mid-log and stationary phase, but was unchanged in the Δ*rbpA*Δ*rbpB* mutant in either growth condition, implying a genetic interaction between *rbpA* and *rbpB* in the regulation of the CPS2 locus. Expression patterns for all mutants were similar between mid-log and stationary phase conditions, except for the CPS7 locus. CPS7 was upregulated in Δ*rbpA* and Δ*rbpA*Δ*rbpB* mutants compared to wild-type in mid-log cells but only the Δ*rbpA*Δ*rbpB* mutant showed a difference with wild-type in stationary phase.

**Figure 3:**
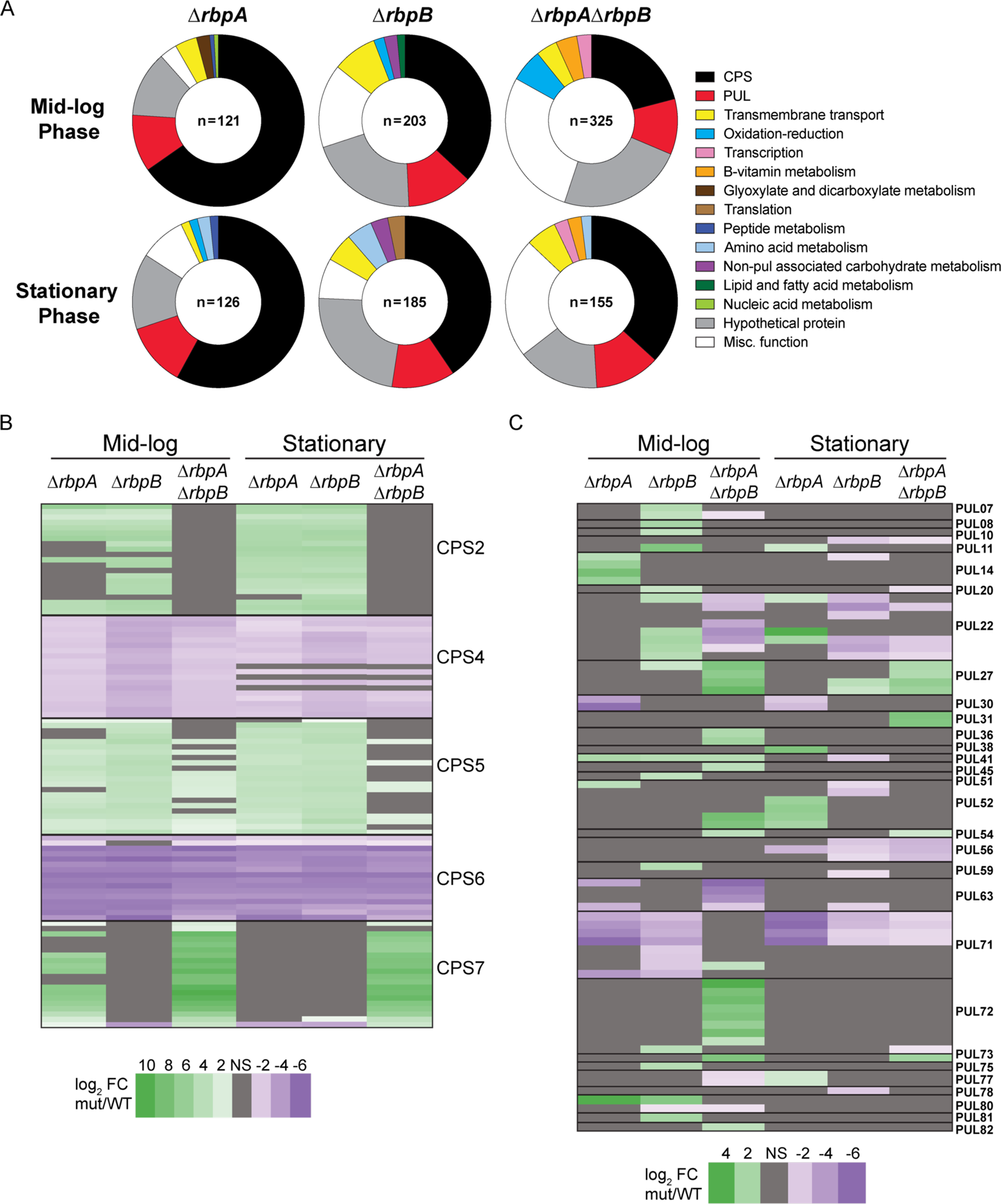
Loss of RBPs leads to altered expression of PULs and CPS loci. (A) Functional categories enriched in differentially regulated genes in rich media RNA-sequencing (genes with a log_2_FC ≥ +1 or ≤ −1 with a q-value <0.06). (B-C) Differentially regulated genes for PULs and CPS loci. Only genes that were significantly differentially regulated are shown. Gene names listed in Dataset S2A.

PULs were the second most abundantly represented functional group among differentially regulated genes (Fig. 3A). Of the 88 annotated PULs (5), 29 (33%) had at least one differentially regulated gene in *rbp* mutant strains compared to wild-type, accounting for 29% (75/263) of genes across the 29 PULs (Fig. 3C). In contrast to CPS expression, PUL expression differences in mutant strains frequently varied according to growth phase. PUL56 was downregulated in all three mutant strains exclusively during stationary phase. In contrast, PUL71 was downregulated in all three mutant strains during stationary phase but in mid-log was only downregulated in single mutants Δ*rbpA* and Δ*rbpB* (Fig. 3C). Similar to the CPS loci, several PULs demonstrated expression patterns indicative of a genetic interaction between *rbpA* and *rbpB*, including PUL22.

PUL22 was upregulated in the Δ*rbpB* mutant but downregulated in Δ*rbpA*Δ*rbpB* double mutant during mid-log growth. In contrast, in stationary phase, PUL22 was upregulated in Δ*rbpA* and downregulated in Δ*rbpB* and Δ*rbpA*Δ*rbpB* mutant strains. Several PULs including PUL08, 10, 14, 51, 59, 75, 80, and 81, were differentially regulated in specific single mutant strains but not differentially regulated in the Δ*rbpA*Δ*rbpB* double mutant. We also saw some expression patterns that may be indicative of redundant regulation by RbpA and RbpB – PULs 36, 45, 54, 72, 73, and 82 were not differentially expressed in the single deletion mutants but were differentially expressed in the Δ*rbpA*Δ*rbpB* double mutant. Collectively, these results suggest that RbpA and RbpB play global roles in *B. thetaiotaomicron* gene expression, and in particular suggest that they coordinate capsular polysaccharide production and carbohydrate utilization through control of CPS and PUL genes, respectively.

### The Δ*rbpB* mutant is defective for growth on raffinose family oligosaccharides

To determine whether *B. thetaiotaomicron* RBPs are required for growth on specific carbohydrates, we carried out an initial screen of Δ*rbpA*, Δ*rbpB*, and Δ*rbpA*Δ*rbpB* strains for growth defects on Biolog plates containing a variety of carbon sources (Fig. S2 and Dataset S3). All three strains were defective for utilization of a number of dietary and host-associated glycans (Fig. S2 and Dataset S3). As observed for the transcriptome, phenotypes for the double mutant Δ*rbpA*Δ*rbpB* strain did not recapitulate all growth defects observed in single mutant Δ*rbpA* or Δ*rbpB* strains, implying a genetic interaction between *rbpA* and *rbpB*. For example, the Δ*rbpA* strain showed faster growth than the wild-type strain on maltotriose, α-methyl-D-galactoside, α-D-lactose, and lactulose while showing slower growth on sucrose, D-trehalose, turanose, D-mannose, and palatinose as compared to wild-type. Defects on turanose, D-trehalose, and palatinose were recapitulated in the Δ*rbpA*Δ*rbpB* strain, but the other growth changes seen in the Δ*rbpA* strain were not observed in the Δ*rbpA*Δ*rbpB* strain. The Δ*rbpB* strain grew slower than wild-type on D-melibiose, β-methyl-D-galactoside, palatinose, and mannan, and defects on β-methyl-D-galactoside, palatinose, and mannan were also observed for the Δ*rbpA*Δ*rbpB* strain. Interestingly, all three strains were defective for growth on palatinose and a methylated galactoside. Unique to Δ*rbpA*Δ*rbpB* was slow growth on gentiobiose and N-acetyl-D-galactosamine. Overall, these results are consistent with transcriptome results that suggest that both *rbpA* and *rbpB* play a role in regulation of carbohydrate utilization.

One of the carbohydrates on which the Δ*rbpB* mutant alone had substantial growth defects was D-melibiose, a subunit of the raffinose family oligosaccharides (RFOs) (Fig. S2). Since RFOs are prevalent in the human diet and are available to organisms that can metabolize them in the distal gut, we chose this phenotype for further evaluation. RFOs consist of the disaccharide sucrose (α-1,2-glucose-fructose) bound to repeating α-1,6-galactosyl residues producing the trisaccharide raffinose and the tetrasaccharide stachyose (Fig. 4A), along with the pentasaccharide verbascose. In addition to sucrose, the α-1,6-galactose-glucose disaccharide melibiose is an RFO subunit. When grown in minimal medium with RFOs or their subunits as the sole carbon source, the Δ*rbpB* strain displayed growth defects on melibiose, raffinose, and stachyose (Fig. 4B). The doubling time of the wild-type strain on minimal medium with melibiose as the sole carbon source was 2.86 hours compared to twice that for the Δ*rbpB* strain (5.76 hours). The Δ*rbpA* and Δ*rbpA*Δ*rbpB* strains did not have growth defects on these substrates, again consistent with possible genetic interactions between RbpA and RbpB with respect to growth on RFOs. Δ*rbpA*, Δ*rbpB*, and Δ*rbpA*Δ*rbpB* strains showed no growth defects on monosaccharide subunits of RFOs, including glucose, galactose, and fructose, or on the disaccharide sucrose (Fig. 4B-C), suggesting that the Δ*rbpB* growth defect is due to the inability of this strain to utilize sugars containing the α-1,6-galactose-glucose linkage.

**Figure 4:**
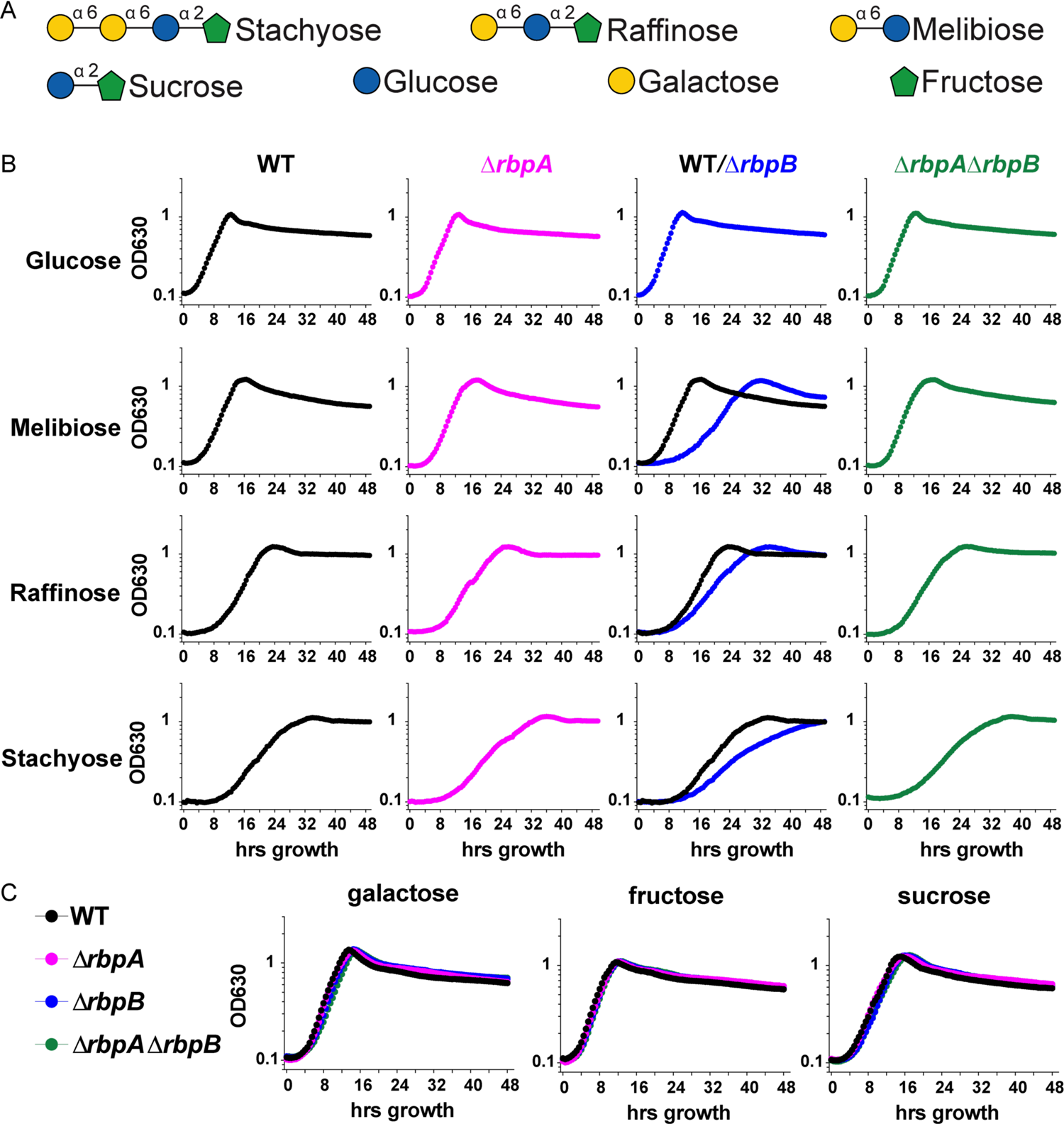
Δ*rbpB* is defective for growth on raffinose family oligosaccharides. (A) Raffinose family oligosaccharides and their subunits. (B-C) Representative normalized growth curves in minimal media from a single biological replicate (n=3) for wild-type (WT) and *rbp* mutants with optical densities (OD_630_) recorded every 30 minutes.

Complementation of the Δ*rbpB* strain was attempted with two different constructs (Fig. S3A). Neither complementation construct restored growth of the Δ*rbpB* mutant on RFOs (Fig. S3B). We measured *rbpB* mRNA levels from wild-type (*rbpB*^+^), Δ*rbpB,* and both complementation strains and found that levels of *rbpB* mRNA in complementation strains were significantly lower than in the wild-type strain (Fig. S3C), which may account for the inability to restore growth on melibiose.

Upon further inspection of the *rbpB* (*BT1887*) native locus in our TYG RNA-seq data, we noticed reduced expression of the immediately adjacent genes *BT1886*, *BT1885*, and *BT1884* in the Δ*rbpB* strain, especially in mid-log phase, suggesting a possible polar effect of the *rbpB* mutation on *BT1886*-*BT1884* (Fig. S4A). We also observed a single transcription start site upstream of *rbpB-BT1884* in TYG (21), a terminator prediction after the *rbpB* ORF, and a terminator prediction after *BT1884* (21), suggesting *rbpB* may be expressed as both a monocistronic mRNA and polycistronic with *BT1886-BT1884* (Fig. S4B). To determine if we could detect *rbpB* co-transcription with *BT1886-BT1884* under conditions relevant to the *rbpB* mutant phenotype we conducted RT-PCR on RNA samples harvested from wild-type (*rbpB*^+^) cells grown to mid-log phase on minimal media with glucose or melibiose. Primer sets spanning junctions between each gene in the putative operon yielded PCR products (Fig. S4C), suggesting that *rbpB* and *BT1886-BT1884* are expressed as an operon. *BT1886*, *BT1885*, and *BT1884* encode a putative RhlE DEAD-box RNA helicase, a hypothetical protein, and a cold shock domain containing protein, respectively. The operon structure suggests the functions of these proteins are linked. One other possibility that may explain the inability to complement the *rbpB* mutant melibiose growth phenotype is that the appropriate stoichiometry of these proteins was not restored by the complementation constructs.

### Loss of RbpB leads to decreased expression of an essential melibiase in PUL24

Given the inability of Δ*rbpB* to utilize α-1,6 linked RFOs, we hypothesized that a gene (or genes) encoding an α-galactosidase would be differentially regulated in the Δ*rbpB* mutant strain compared to wild-type. Although there are several α-galactosidases annotated in the genome (7), none of them were significantly differentially regulated in our TYG RNA-seq, suggesting differential regulation may be specific to growth in minimal medium with melibiose. We therefore performed more RNA-seq to identify candidate genes responsible for this phenotype. We compared transcriptome profiles of wild-type and Δ*rbpB* strains grown in minimal media with glucose or melibiose. To identify genes that are uniquely transcriptionally responsive to the α-1,6 linkage in melibiose, we also compared the glucose- and melibiose-grown cells’ transcriptomes to that of cells grown in minimal medium with a 1:1 mixture of glucose and galactose, the monosaccharides that makeup melibiose.

Comparing wild-type and Δ*rbpB* transcriptomes in all three media, we identified genes in PUL24 that were strongly differentially regulated (Fig. 5A and Dataset S2B). PUL24 (genes *BT1871-BT1878*) contains a SusC/D-like pair (BT1874-BT1875), a σ/anti-σ factor pair (BT1876-BT1877), and four putative glycosyl hydrolases belonging to families GH3 (BT1872), GH43 (BT1873), GH76 (BT1878), and GH97 (BT1871).

**Figure 5:**
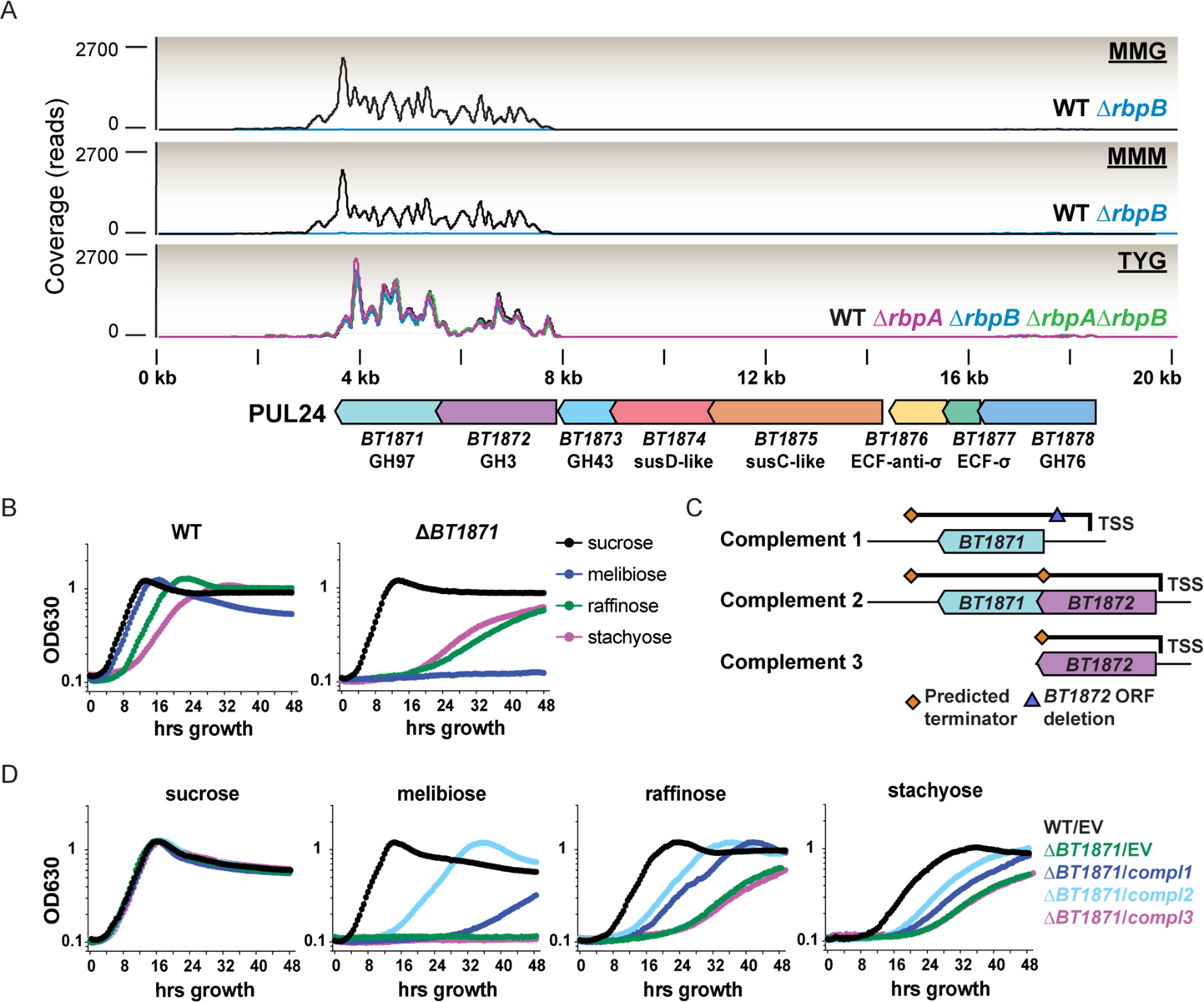
Loss of *rbpB* leads to loss of expression of an essential melibiase in PUL24. (A) Normalized expression coverage curves for mid-log phase cultures in PUL24 with putative gene annotations. (B and D) Representative growth curves in minimal media from a single biological replicate (n=3) with optical densities (OD_630_) recorded every 30 minutes. (C) Genomic regions inserted into pNBU2 vectors for complementation of *BT1871-BT1872* shown in (D). (D) Empty vector (EV) controls contain integrated pNBU2 without an insert.

Genes *BT1873-BT1878* were expressed at very low levels in wild-type and Δ*rbpB* strains in all of the conditions we tested (Fig. 5A, Dataset S2B) suggesting that these genes are not involved in glucose, galactose, or melibiose metabolism. However, *BT1871* and *BT1872* were highly expressed in the wild-type strain growing in glucose and melibiose (Fig. 5A) (and the glucose-galactose mixture, Dataset S2B) but were expressed at barely detectable levels in the Δ*rbpB* strain. In contrast, *BT1871* and *BT1872* were not differentially expressed between wild-type and Δ*rbpB* strains grown in TYG (Dataset S2A).

Previous work showed that BT1871 has *in vitro* melibiase activity (61), and transposon insertions in *BT1871* led to decreased fitness in melibiose in a carbohydrate utilization screen (62). To confirm that BT1871 was important for *B. thetaiotaomicron* utilization of RFOs including melibiose as a sole carbon source, we deleted *BT1871* and cultured the Δ*BT1871* and wild-type strains in minimal media with melibiose, raffinose, stachyose, and sucrose (Fig. 5B). The Δ*BT1871* strain showed no growth defect on sucrose compared to the wild-type strain, which is expected based on its predicted melibiase activity. In contrast, the Δ*BT1871* mutant could not grow on melibiose, indicating *BT1871* is essential for melibiose utilization. Additionally, the Δ*BT1871* mutant showed reduced growth on raffinose and stachyose compared to wild-type (Fig. 5B), indicating that *BT1871* is required for metabolism of RFOs in general. Residual growth of the Δ*BT1871* mutant on raffinose and stachyose is presumably due to the ability to utilize fructose from the α-1,2 sucrose linkage.

We constructed three different complementation strains to confirm that BT1871 is responsible for the melibiose growth defect. There is a single predicted promoter upstream of *BT1872* (21) and the *BT1872* and *BT1871* open reading frames are separated by only 32 bp suggesting that they are co-expressed. Complementation strain 1 (compl1, Fig. 5C) carried the native promoter upstream of *BT1872* followed by a deletion of the *BT1872* ORF and the intact *BT1871* gene. Complementation strain 2 (compl2, Fig. 5C) carried the intact promoter and *BT1872* and *BT1871* genes.

Complementation strain 3 (compl3, Fig. 5C) carried the promoter and *BT1872* only. The compl1 construct partially restored growth on melibiose, raffinose, and stachyose (Fig. 5D). The compl2 construct improved growth on RFOs compared to the compl1 construct, whereas the compl3 construct (*BT1872* alone) failed to complement (Fig. 5D) Taken together these results indicate that *BT1871* is an essential melibiase required for RFO utilization and the decrease in *BT1871* mRNA in the Δ*rbpB* strain is responsible for the melibiose growth defect.

## Discussion

Though it is well established that rapid nutrient shifts affect the composition and metabolic activities of gut microbes (47, 63–65), the regulatory mechanisms that allow them to sense and rapidly adapt to use of different nutrient sources are poorly understood. Canonical mechanisms for global transcriptional regulation of carbon source utilization in model organisms from the phyla Proteobacteria and Firmicutes(66) are absent in the Bacteroidetes (67–69). Likewise, RNA chaperones and RNA-mediated post-transcriptional regulatory mechanisms that coordinate metabolism and responses to changing environmental conditions (28, 70–73) are commonly found in Proteobacteria and Firmicutes but have not been described in Bacteroidetes. In this study, we identify a family of conserved RNA-binding proteins that is broadly distributed among members of the Bacteroidetes and some Proteobacteria that lack canonical RNA chaperone regulators. These RBPs occur in multiple copies in a given genome and are highly expressed in a number of *Bacteroides* species from the human gut in culture and in mouse models. At least one of these proteins from *B. thetaiotaomicron*, RbpB, is able to bind ssRNA *in vitro* in a sequence specific manner. Deletion of *rbpA* and *rbpB* in *B. thetaiotaomicron* leads to global dysregulation of CPS loci and PULs and perturbed growth on a variety of carbohydrate sources. Overall, these results suggest that this family of RBPs may play global regulatory roles in carbohydrate metabolism in the *Bacteroides*.

While our study provides strong evidence for the importance of these RBPs in global regulation of gene expression, the mechanisms by which RBPs mediate these effects are still unknown. We hypothesize that RBPs act as RNA chaperones that control mRNA stability and translation by binding to target mRNAs and modulating ribosome association or access of RNases (Fig. 6). RBP modulation of mRNA translation or stability may be through direct interaction of RBPs with target mRNAs (Fig. 6A) or through facilitating base pairing of sRNAs to mRNAs (Fig. 6B), either of which could result in changes to mRNA structure that alter accessibility to ribosomes or RNases. Little is known about RNA chaperone function in the Bacteroidetes, but in the case of RbpB, its role in RNA metabolism is supported by its genomic location.

**Figure 6:**
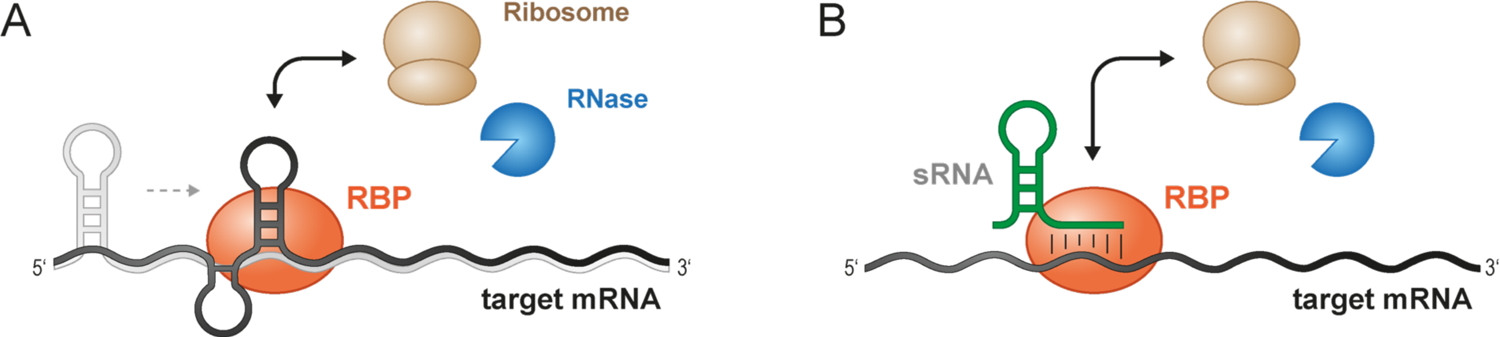
Model for possible mechanisms of RBP-mediated regulation (A) RBP binding directly to mRNAs could promote structural changes (represented by the transition from light gray to dark gray conformation) that alters access of ribosomes or RNases to change translation or mRNA stability. (B) RBPs could facilitate sRNA binding to mRNA targets and alter access of ribosomes or RNases to change translation or mRNA stability.

Annotations for BT1885 (DEAD-box RNA helicase) and BT1884 (cold-shock protein) suggest that *rbpB-BT1884* may be an RNA metabolism operon. To date, we do not have evidence supporting or refuting a role for RBPs in modulation of sRNA function. However, recent literature suggests that sRNAs may play an important role in modulation of carbohydrate metabolism in *Bacteroides* (19, 21). One recent study described an N-acetyl-D-glucosamine-inducible sRNA called GibS that binds *in vitro* to mRNAs involved in carbohydrate metabolism. Mutant strains lacking GibS had nine differentially-regulated genes compared to the wild-type parent strain. Two of these were *BT1871-BT1872,* where expression was reduced in the Δ*gibS* compared to the wild-type strain. GibS binding to *BT1871* mRNA was predicted *in silico* but could not be demonstrated *in vitro*. The authors speculated that GibS binding to *BT1871* mRNA required an unidentified RNA chaperone. To test whether RbpB facilitates RFO utilization by a GibS-dependent mechanism, we generated deletion mutants Δ*gibS* and Δ*rbpB*Δ*gibS* and grew these strains alongside wild-type in the presence of melibiose (Fig. S5A). The Δ*gibS* mutant grew similarly to wild-type and the Δ*rbpB*Δ*gibS* mutant grew similarly to the Δ*rbpB* parent strain on melibiose (Fig. S5A). RT-qPCR showed that levels of *BT1871* mRNA were also similar between wild-type and Δ*gibS* strains (Fig. S5B), suggesting that GibS does not play a major role in modulating *BT1871* mRNA levels in our growth conditions. Overall, these results suggest that under our growth conditions, RbpB regulates *BT1871* independently of GibS.

We have yet to explore the role of RBPs in helping *B. thetaiotaomicron* colonize or be maintained in the host gut. In a recent study (62) screening transposon (Tn) mutants for a wide variety of *in vitro* and *in vivo* phenotypes, there were no reported insertions in *rbpA* or *rbpB*. Insertions in *rbpC* led to reduced fitness on glucose-containing media (62), possibly explaining our inability to generate Δ*rbpC* mutants in our standard glucose-rich media. The *rbpC* mutants also had an increased growth on melibiose suggesting that *rbpC* also plays a role in utilization of RFOs. In the same study (62), colonization of germ-free mice fed a plant polysaccharide-rich diet with the *B. thetaiotaomicron* Tn-mutant pool led to increased fitness of *rbpC* mutants. *BT1871* mutants showed decreased fitness over time, whereas Tn-insertions into several other PUL24 genes led to increased fitness *in vivo*. Overall, these data indicate that RBPs and the PULs they regulate may be important for *in vivo* fitness.

Regulation by RBPs may represent a critical mechanism for coordination of carbohydrate utilization and production of cell surface capsular polysaccharides. Differential regulation of PULs and CPS loci in *rbp* deletion strains is consistent with several reports indicating a regulatory link between these polysaccharide metabolic processes in *B. thetaiotaomicron* (17, 74, 75). Our RNA-seq data showed that deletion of RBPs leads to reduced expression of CSP4 and CPS6 and increased expression of CPS5. In *B. thetaiotaomicron,* CPS4 is normally the most highly expressed locus *in vitro* and in mouse models when dietary glycans are present (64, 75, 76), and disruption of CPS4 expression leads to decreased fitness in mouse competitions (77, 78). In a study monitoring CPS expression in a mouse model over time, it was observed that even when the *B. thetaiotaomicron* inoculum expressed one dominant CPS locus, expression over time varied between mice (75). While CPS4 was most often highly expressed in mice fed a high-fiber diet, mice on fiber-free diets typically expressed CPS5 or CPS6. *B. thetaiotaomicron* mutants that could only express a single CPS locus had a decreased ability to recover from antibiotic-induced stress (75). These studies along with our present work collectively suggest that the ability to shift among different CPS types is advantageous in the host and that this regulation may be mediated in part by RBPs. Further characterization of the RBPs and their regulatory mechanisms may provide critical insight into how *Bacteroides* coordinately control carbohydrate availability with cell surface properties. This could reveal key principles governing mechanisms in host dynamics.

## Materials and Methods

### Bacterial culturing and genetic manipulation

*B. thetaiotaomicron* VPI-5482 strains were grown anaerobically in a Coy Laboratory Products vinyl anaerobic chamber with an input gas of 20% CO_2_, 10% H_2_, 70% N_2_ balance. Routine culturing of *B. thetaiotaomicron* was done in Tryptone-Yeast-Extract-Glucose (TYG) (79) broth and on Difco Brain Heart Infusion (BHI) agar plates with 10% defibrinated Horse Blood (HB) (Quad Five) at 37°C. *Escherichia coli* strains were grown aerobically at 37°C on BHI-10%HB for conjugations and Luria Broth for all other applications. Minimal medium (22) was supplemented with B_12_ [3.75nM final] (Sigma) and carbohydrates as needed at the following final w/v concentrations unless otherwise indicated: 4.0% stachyose (Sigma), 2.0% D-(+)-raffinose (Sigma), 0.5% D-(+)-melibiose (Sigma), 0.5% α-D-glucose (Sigma), 0.5% D-(+)-galactose (Sigma), 0.5% β-D-(-)-fructose (MP Biomedicals), and 0.5% sucrose (MP Biomedicals). When needed, antibiotics were added at the following final concentrations: 100 µg/ml ampicillin (Sigma), 200 µg/ml gentamicin (Goldbio), 25 µg/ml erythromycin (VWR), 200µg/ml 5’-fluoro-2’-deoxyuridine (VWR), 100ng/ml anhydrotetracycline (Sigma), 25 µg/ml kanamycin (Fisher). All strains, vectors, and primers are listed in Table S1. For all experiments, wild-type *B. thetaiotaomicron* is Δ*tdk* (strain AA0014 in Table S1).

Markerless deletions were made in *B. thetaiotaomicron* using the pExchange_*bla*_*tdk*_*ermGb* (80) and the pLGB13*_bla_ermG* (81) suicide vector-based allelic exchange methods. Upstream and downstream regions of the gene to be deleted were amplified using Kappa HiFi (Kappa Biosystems) and cloned into pExchange_*bla*_*tdk*_*ermGb* using standard restriction digest and ligation methods and splicing by overlap exchange (SOE) (22). Alternatively, inserts were cloned into Q5 (NEB) amplified pExchange_*bla*_*tdk*_*ermGb* using restriction digest and ligation of a gBlock insert (IDT). GibS flanks were Q5 amplified and ligated to restriction-digested pLGB13*_bla_ermG.* Complete vectors were conjugated into *B. thetaiotaomicron* with *E. coli* S17 λ-pir using established methods (22). Complementation pNBU2_*bla*_*ermGb* vectors (5, 82) were cloned using standard restriction digest and ligation methods and conjugated into *B. thetaiotaomicron* as done with pExchange. pNBU2_*bla*_*ermGb* vectors were PCR screened for insertion into a single attachment site as done previously (5). The pET-28a-*rbpB* protein expression vector was generated by inserting the *rbpB* (*BT1887*) ORF 5’ to the thrombin cleavage site and 6xHis tag in the pET28a backbone. pET-28a-*rbpB* was cloned using Q5 PCR amplification and NEBuilder assembly (NEB) in *E. coli* XL10-Gold competent cells (Agilent) before being moved into *E. coli* BL21 (DE3) for protein expression.

### Computational identification of RNA regulators in human gut-associated microbial genomes

To identify genomes containing CsrA, ProQ, RRM-1, and Hfq in the human gut microbiome, we utilized a custom database of 313 human gut-associated microbial genomes containing a single representative genome for a species (22, 37). Candidate RNA regulator genes were identified using hmmer v3.3 (hmmer.org) with trusted cutoffs and the individual hidden Markov model from each protein queried: Hfq, PF17209.4; CsrA, PF02599.17; ProQ, PF04352.14; RRM-1, PF00076.23 (39). The resulting gene list was then run against Pfam-A.hmm version 33.1 using hmmer to verify that the query PFAM was the top hit for the target domain using trusted cutoff values. ProQ PF04352.14 gene hits that also contained an N-terminal FinO_N domain (PF12602.9) were removed from the final annotation list. RRM-1 PF00076.23 gene hit list was limited to less than 150 amino acids to remove a few genes containing transmembrane domains. The maximum likelihood phylogenetic tree in Fig. 1A was built using a multisequence alignment of 13 conserved core genes (AspS, Ffh, FusA, GltX, InfB, LeuS, RplB, RpsE, RpsH, RpsK, TopA, TufA, RpoB) identified and described previously (83). Briefly, protein sequences for each group of orthologs were individually aligned with MUSCLE (84), concatenated, and subjected to phylogenetic reconstruction with RAxML(85). The phylogeny was visualized using FigTree (http://tree.bio.ed.ac.uk/software/figtree/).

### RBP expression in publicly available RNA-seq datasets

Publicly available RNA-seq datasets were downloaded from NCBI (see Table S2) for sample IDs. RNA-seq reads were quality filtered with Trimmomatic v0.36 (86). Read mapping and sample normalization was calculated with Rockhopper v2.03 (87, 88) and normalized expression values were graphed using JMP v15 (89).

### RbpB EMSAs and motif identification

#### Purification of RbpB

*E. coli* BL21 (DE3) cells with the pET-*rbpB* vector were grown to late exponential phase (0.6-0.8 OD_600_ as measured on an Ultraspec 2100 Pro, Amersham) and protein expression induced with 1 mM final IPTG (Goldbio) for 4 hr at 37 °C. Cells were harvested by centrifugation and pellets resuspended in 30 ml of extraction buffer (1X PBS, 0.5M NaCl, pH 7.2) before being lysed in a French press. Supernatant was collected after centrifugation at 16,000 × g for 10 min at 4°C. The supernatant was then fractionated using a HiTrap Ni^2+^ column (GE Healthcare) following the manufacturer’s instructions. Fractions containing RbpB were dialyzed overnight in TGED buffer (10 mM Tris-HCl pH 8, 5% glycerol, 0.1mM EDTA, and DTT 0.015 mg/mL) and loaded onto a HiTrap-Q column (GE Healthcare). The column was washed with TGED buffer, and protein was eluted with a linear gradient of NaCl (0.1 M to 1 M) in TGED buffer. The fractions containing the protein were pooled, dialyzed, and concentrated using Centricon 10 concentrators (Millipore-Sigma), mixed with an equal volume of 100% glycerol and stored at −20°C.

#### Radiolabeled pentaprobe synthesis

Twelve pentaprobes containing all the possible 5-nt combinations were prepared based on a published protocol with some modifications (52). All oligonucleotides used in pentaprobe synthesis are listed in Table S1. Single-stranded oligonucleotides for PP1-PP6 were Q5 PCR amplified with a 5’ T7 promoter for either the Watson strand or the Crick strand, generating twelve dsDNA templates with a single T7 site, two each for PP1-PP6. The Watson strand of PP1 dsDNA is identical to the coding strand of the PP1 pentaprobe, and the Crick strand is identical to the coding strand of the PP7 pentaprobe. These dsDNA fragments were then used to produce twelve different ssRNA pentaprobes by *in vitro* transcription from the T7 promoter with a MEGAscript T7 kit (Ambion). Transcribed RNA fragments were 5ʹend-labeled with ATP [γ^32^P] (PerkinElmer) using the KinaseMax kit (Ambion) following the manufacturer’s protocol.

#### Electrophoretic mobility shift assay and motif prediction

RNA-protein gel electrophoretic mobility shift assays were performed using 0.01 pmol of ^32^P-labeled pentaprobe RNA and the indicated amounts of RbpB in binding buffer (10mM Tris-HCl pH 8.0, 0.5 mM DTT, 0.5 mM MgCl_2_, 10mM KCl, 5mM Na_2_HPO_4_–NaH_2_PO_4_ pH 8.0). The mixture was incubated at 37 °C for 30 minutes, and non-denaturing loading buffer (50% glycerol and 0.1% bromophenol blue) was added. The samples were resolved on a 4.6% native polyacrylamide gel for 1.5 hours at 10 mA. The fraction of RbpB bound was determined using a Fluorescent Image Analyzer FLA-3000 (FUJIFILM) to quantify the band intensities. *K*_D_ values were calculated using Sigmaplot software based on a published method (90). The MEME program (55) was used to predict conserved motifs for the positive pentaprobe sequences with the following parameters: maximum number of motifs, 10; minimum motif width, 4; and maximum motif width, 8. *K_D_* was calculated for RbpB binding to PP3 and motif 1 using three technical replicates as done previously (56).

### RNA sequencing sample prep and processing

For rich media RNA-seq, strains were cultured in 5 ml of TYG in biological triplicate to stationary phase overnight. Each culture was then sub-cultured 1:100 into two 5 ml TYG cultures. One tube was cultured to mid-log (0.35-0.6 OD_600_) and the second tube was cultured to early stationary phase (1.2-1.4 OD_600_) as measured in a Thermo Spectronic 200 (Thermo Fisher Scientific brand, referred to as ThermoSpec below). Then 500 μl of cells were then spun down at 7,500 x g for 3 min at room temperature, supernatant removed, and pellets re-suspended in 600 μl of TriReagent (Sigma). RNA was then isolated from the re-suspensions using the Zymo Direct-Zol RNA Mini-Prep kit (Zymo) which includes on-column DNaseI treatment. RNA quality was evaluated using a Qubit 2.0 fluorometer (Invitrogen) and Agilent 2100 Bioanalyzer (UIUC Biotechnology Center). Total RNA was then submitted to the W. M. Keck Center for Comparative and Functional Genomics at UIUC for rRNA depletion, library construction, and sequencing. Briefly, ribosomal RNAs were removed from total RNA with an Illumina Ribo-Zero rRNA Removal Kit for Bacteria. RNA-seq libraries were produced with a ScriptSeq v2 kit (Illumina) and cleaned with AMPure beads (Beckman Coulter) to remove any fragments <80 nt. Libraries were sequenced on an Illumina HiSeq2500 using HiSeq SBS sequencing kit v4 to give 160 nt single-end reads. Results were de-multiplexed with bcl2fastq v2.17.1.14 (Illumina). Reads were quality filtered and trimmed using Bioconductor package ShortRead (91) to first remove reads with >1 N or if ≥75% of a read is a single nucleotide and then the first 2 nucleotides were removed from each sequence read. Sequencing adapters were removed with fastx_clipper (http://hannonlab.cshl.edu/fastx_toolkit/index.html). Residual rRNAs were removed using bowtie2 (92, 93) and final reads mapped to the genome and analyzed using Rockhopper v2.03. All raw Rockhopper calculated expression values were increased by 1 and fold changes (FC) calculated as log_2_(mutant expression value + 1/wild-type expression value +1). RNA-seq processing statistics are summarized in Dataset S2C.

For RNA-seq of cultures grown in minimal media, a single colony/strain was smeared onto half a 100 mm BHI-10%HB agar plate with a cotton swab and cultured for 24 hours. Lawns were then re-suspended in 5 ml of minimal medium + glucose (MMG) and spun down at 4000g x 5 min. Cell pellets were washed three times with 1 ml of MMG and then diluted to 0.07 OD_630_ in 200 μl of MMG, as measured on a Biotek Synergy HT plate reader (referred to as Biotek below). Cells were then diluted 1:1000 in 25 ml of MMG and cultured overnight to 0.35-0.50 OD_600_ (ThermoSpec). Cells were then spun down and re-suspended in 1 ml of minimal medium without a carbon source per every 5 ml of culture. For each biological replicate, these suspensions were then diluted to 0.1 OD_600_ (ThermoSpec) in 5 ml of MMG, minimal medium + melibiose (MMM), or minimal medium + 0.25% w/v glucose + 0.25% w/v galactose (MMGG).

Cultures were then grown to 0.45-0.65 OD_600_ (ThermoSpec), and then stabilized in Qiagen RNA protect. Briefly, 4 ml of culture was combined with 8 ml of RNA protect, vortexed, and then incubated at room temperature for 5 min. Suspensions were then spun at 4,000 x g for 10 minutes at 4°C. Supernatants were decanted and pellets stored at −80 °C. Cell pellets were thawed, and RNA prepped using a Qiagen RNeasy Mini Kit. RNA was sequenced and analyzed as done above with the RNA-seq for cells grown in TYG, with the exception that libraries were sequenced on an Illumina HiSeq4000 to produce 150 nt single-end reads.

### RNA-seq protein functional and pathway analyses

RNA-seq data from each mutant strain (Δ*rbpA*, Δ*rbpB*, Δ*rbpA*Δ*rbpB*) was compared to wild-type in rich medium at mid-log and stationary phase, yielding six comparisons. All genes with a log_2_FC ≥ +1 or ≤-1 with a q-value <0.06 were assigned to functional groups using the following framework. GSEA was used on each of the six differentially expressed gene sets individually to identify enriched functional groups. *B. thetaiotaomicron*-specific functional gene sets used included: KEGG pathways, CPS loci, PULs, and corrinoid transport (22). GSEA identified enriched gene sets corresponding to PULs, CPS loci, TCA cycle (KEGG pathway bth00020), corrinoid transport, and microbial metabolism in diverse environments (bth01120). Since these were identified as enriched categories, if one of the significantly differentially regulated genes in Dataset S2A was part of one of these gene sets, it was assigned that as a category identifier with the exception of “microbial metabolism in diverse environments.” GSEA-identified enriched genes in this KEGG category were further split into sub-categories including TCA cycle and glyoxylate and dicarboxylate metabolism (bth00630). If genes were associated with both TCA cycle and glyoxylate and dicarboxylate metabolism, they were assigned TCA cycle since this category was identified as enriched by GSEA directly, but glyoxylate and dicarboxylate metabolism was not. Since corrinoid transport was enriched in our differentially expressed genes, any gene associated with B_12_ metabolism that was in our differentially regulated genes was assigned a “B-vitamin metabolism” category. Remaining gene functions were assigned using gene ontology (GO) from QuickGO. Briefly, gene names were used to extract UniprotKB identifiers that were then used to pull GO biological process annotations from QuickGO when available. Only the first reported GO assignment was used for each gene. GO terms were further grouped into custom functional categories to make a more tractably sized list of functional categories for visualization. Genes without any of the above functional assignments were labeled as either “hypothetical protein” if they were annotated as such or “miscellaneous” if the gene had a putative functional annotation that was not captured by the other functional categories. All resulting functional groups are listed in Dataset S2A.

### Biolog carbon utilization assays

Biolog carbon utilization assays were conducted according to manufacturer recommendations as follows. Biolog carbon source PM1 and PM2A MicroPlates were brought to room temperature to avoid condensation prior to opening the seals. Plates were then cycled into the anaerobic chamber and maintained in an anaerobic desiccant box for 24 hours. Single colonies of each strain were swabbed onto BHI-10% horse blood plates and cultured overnight to produce a lawn of cells. Cells were aerobically suspended into 5 ml of reduced minimal medium without a carbon source to 40% turbidity (OD_590_, ThermoSpec) using a cotton swab. Suspensions were cycled into the chamber and 1.5 ml combined with 22 ml of anoxic, reduced minimal medium without a carbon source. Each carbon source plate was then inoculated with 100 μl of diluted cells and statically incubated for 30 minutes at room temperature to facilitate compound dissolution before measuring time point zero. Plates were statically incubated at 37 °C with manual OD_630_ readings taken every hour in the plate reader for the first 11 hours of growth. Plates were then left in the chamber overnight and optical density readings were resumed after 24 hours of growth. Time points were then taken every 3 hours to a final time point of 36 hours of growth. Linear regression and prediction curves were calculated using Prism. Negative control wells and Xylitol (PM2A) were removed from linear regression calculations. Xylitol was removed due to an unknown occlusion (potentially condensation or precipitation) causing transiently high OD_630_ readings. In the absence of these transient values, *B. thetaiotaomicron* could not grow on Xylitol as a sole carbon source in these experiments.

### Minimal media growth assays

Strains were cultured from a colony in 5 ml of TYG for 24 hours and then sub-cultured 1:1000 into 5 ml of MMG for 24 hours. 1 ml of stationary phase MMG cultures were spun down 4000 g x 10 min at room temperature. Supernatants were removed and pellets resuspended in 1 ml of minimal medium without a carbon source. 2 μl of cells were then sub-cultured into 198 μl of minimal media containing carbon sources to appropriate final concentrations in flat bottom, 96 well Corning Costar tissue culture-treated plates (Sigma). Plates were sealed with a Breathe-Easy gas permeable membrane (Sigma) and statically cultured in the Biotek plate reader for 48 hours with optical density recorded every 30 minutes.

### qRT-PCR of *rbpB* and *BT1871*

Strains were cultured in MMG to stationary phase overnight, sub-cultured 1:100 into 5 ml of MMG, and then cultured to mid-log (0.38-0.52 OD_600_, ThermoSpec). All cultures for strains containing pNBU2_*ermGb* vectors contained erythromycin. Four ml of cells were pelleted at 4,000 g x 10 min, supernatant decanted, and then RNA isolated with a Qiagen RNeasy mini kit. Residual DNA was degraded on-column using Qiagen RNase-Free DNase Set and the RNA cleaned with a Qiagen RNeasy Mini kit. First-strand cDNA synthesis was done with a SuperScript II RT kit (Invitrogen) and random hexamers (Invitrogen). Post reverse transcription, the SuperScript reaction was incubated with 27 μl of 1N NaOH at 65 °C for 30 min, neutralized with 27 μl of 1N HCl, and cleaned up with a Qiagen MinElute PCR purification kit. cDNA was diluted and *rbpB*, *BT1871*, and 16s rRNA copies amplified using 2x QX200 ddPCR EvaGreen Supermix (Biorad) and quantified using the QX200 Droplet Digital PCR system (Biorad) according to manufacturer instructions. All ddPCR consumables were supplied by Biorad and Rainin (pipette tips only). Relative ratios were calculated by dividing the *rbpB* or *BT1871* counts by the 16s rRNA counts.

### Determination of operon structure of *rbpB* in *B. thetaiotaomicron*

Strains were cultured in MMG to stationary phase overnight and then sub-cultured 1:1,000 into 4 ml of MMM and 1:10,000 into 4 ml MMG. MMG and MMM cultures were grown to mid-log (OD_630_ 0.25-0.35, Biotek), pelleted at 4,000g x 10 min at 4°C, and supernatant removed. RNA and cDNA were prepped as done for qPCR with the exception that residual DNA was degraded on beads using an Ambion nuclease-free DNAse kit. Overlap end-point PCR was done with KAPA HiFi (KAPA Biosystems). gDNA was prepped using a Qiagen DNeasy Blood and tissue kit.

## Data Availability

All RNA-seq datasets corresponding to the samples listed in Dataset S2C are publicly available on NCBI under BioProject accession number PRJNA723047.

## Acknowledgements

We thank Alvaro Hernandez, Chris Wright, and staff of the Roy J. Carver Biotechnology Center for assistance with RNA-seq. We also thank Danielle Campbell for guidance on comparative genomics and Auroni Gupta and Saika Hossain for assistance with EMSAs. We are grateful to Sandy Pernitzch (Scigraphix) for assistance with model graphics. This work was funded by UIUC and the UIUC Department of Microbiology, a Roy J. Carver Charitable Trust award (15–4501) to P.H.D. and initial complement funding to P.H.D. from UCR. A.N.D.A. was funded by an Alice Helm Graduate Research Excellence Fellowship and the UIUC Department of Microbiology.

A.N.D.A, M.S.A, Z.A.C., P.H.D, and C.K.V. designed research and analyzed data. A.N.D.A., M.S.A., Z.A.C., X.M., and P.H.D. performed research. A.N.D.A., M.S.A., P.H.D., and C.K.V. wrote the paper.

