## Supplemental Figs and Tables for "A Novel Family of RNA-Binding Proteins Regulate Polysaccharide Metabolism in *Bacteroides thetaiotaomicron*"

### Supplemental materials

### Dataset descriptions

**Dataset S1: Conservation of RNA regulators in the human gut microbiome.**
Table lists the microbial strains and their corresponding Refseq IDs used in Figure 1. Unique abbreviations (Abbv) were assigned to each Refseq strain and used as prefixes for each gene name. IDs following the abbreviation prefix correspond to publicly available gene IDs for each Refseq entry. Gene names are listed for each RNA binding protein identified, with each RRM-1 gene cluster ID labeled in parentheses for copies *rbpA-J*. *rbpA-rbpA'* gene copies not distinguished.

**Dataset S2: RNA-sequencing expression data for TYG and minimal media.**
Rockhopper RNA-seq analysis for significantly differentially regulated genes. Expression (Expr) values represent the raw Rockhopper calculated expression values +1. Rockhopper expression values shown represent the composite of three biological replicates and q-values are calculated by Rockhopper. (A) RNA-seq results for TYG in mid-log (ML) and stationary phase (Stat). Genes shown had at least one condition in which the calculated $log_2FC$ was $\geq +1$ or $\leq -1$ with a q-value $<0.06$. (B) RNA-seq results for minimal media glucose (MMG), minimal media melibiose (MMM), or minimal media glucose-galactose (MMGG). Genes shown had at least one wild-type (WT) to mutant comparison in which the q-value was $<0.06$. (C) RNA-seq mapping statistic, sample IDs, and SRA accession IDs.

**Dataset S3: Biolog carbon sources and optical densities.** Biolog carbohydrates listed with raw $OD_{630}$ readings for 11 and 24 hours of growth for WT, Δ*rbpA*, Δ*rbpB*, Δ*rbpA*Δ*rbpB*.

### Supplemental figures

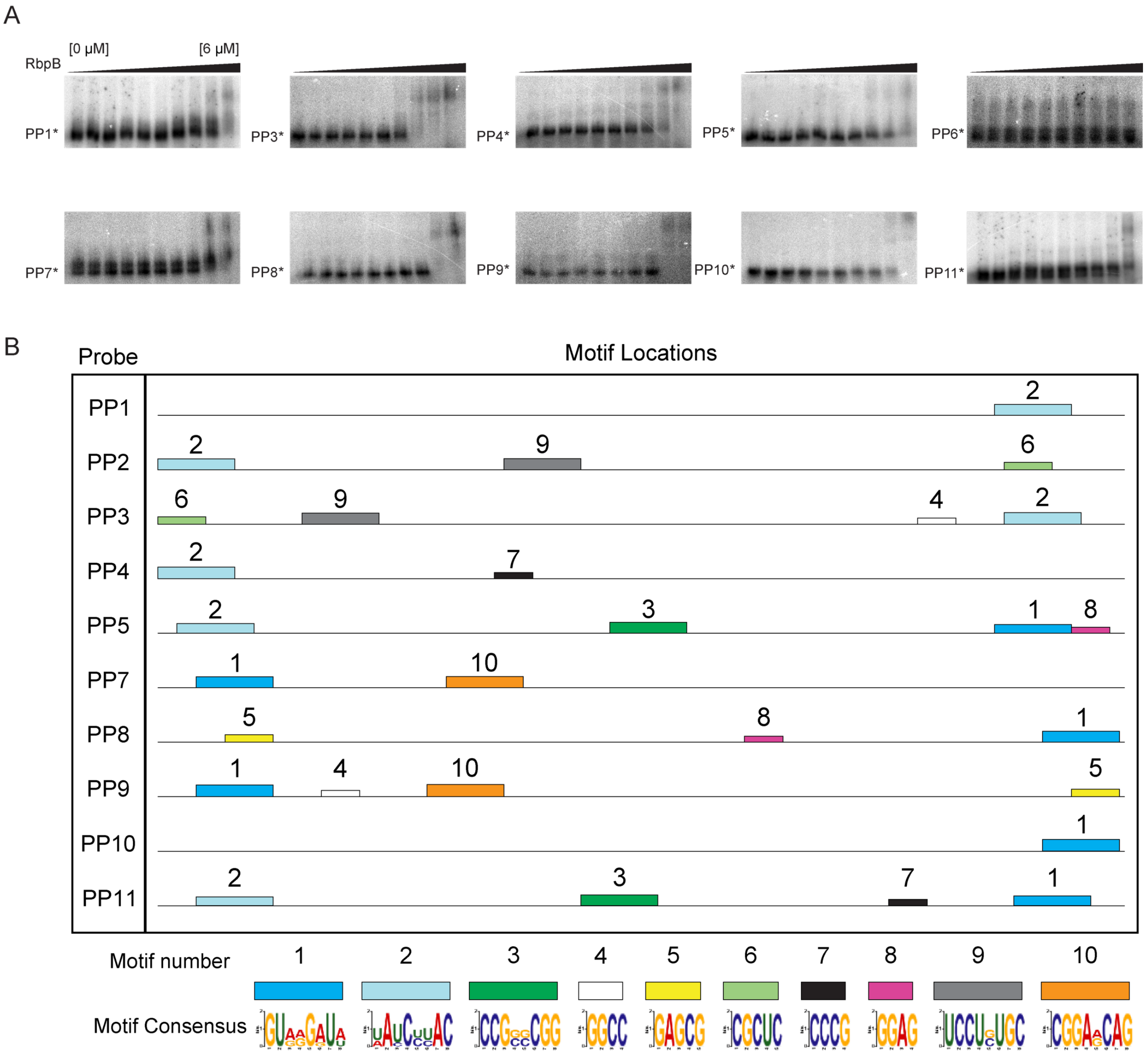


**Figure S1: Candidate RbpB binding motifs identified by MEME.** (A) RbpB-pentaprobe EMSAs for pentaprobes 1 and 3-11 (PP1 and PP3-11). RbpB [μM] increases from left to right as follows: 0, 0.02, 0.05, 0.09, 0.19, 0.38, 0.75, 1.50, 3.00, 6.00. Asterisk * indicates the unbound radiolabeled probe. (B) MEME identified RNA motifs in pentaprobes 1-5 and 7-11.

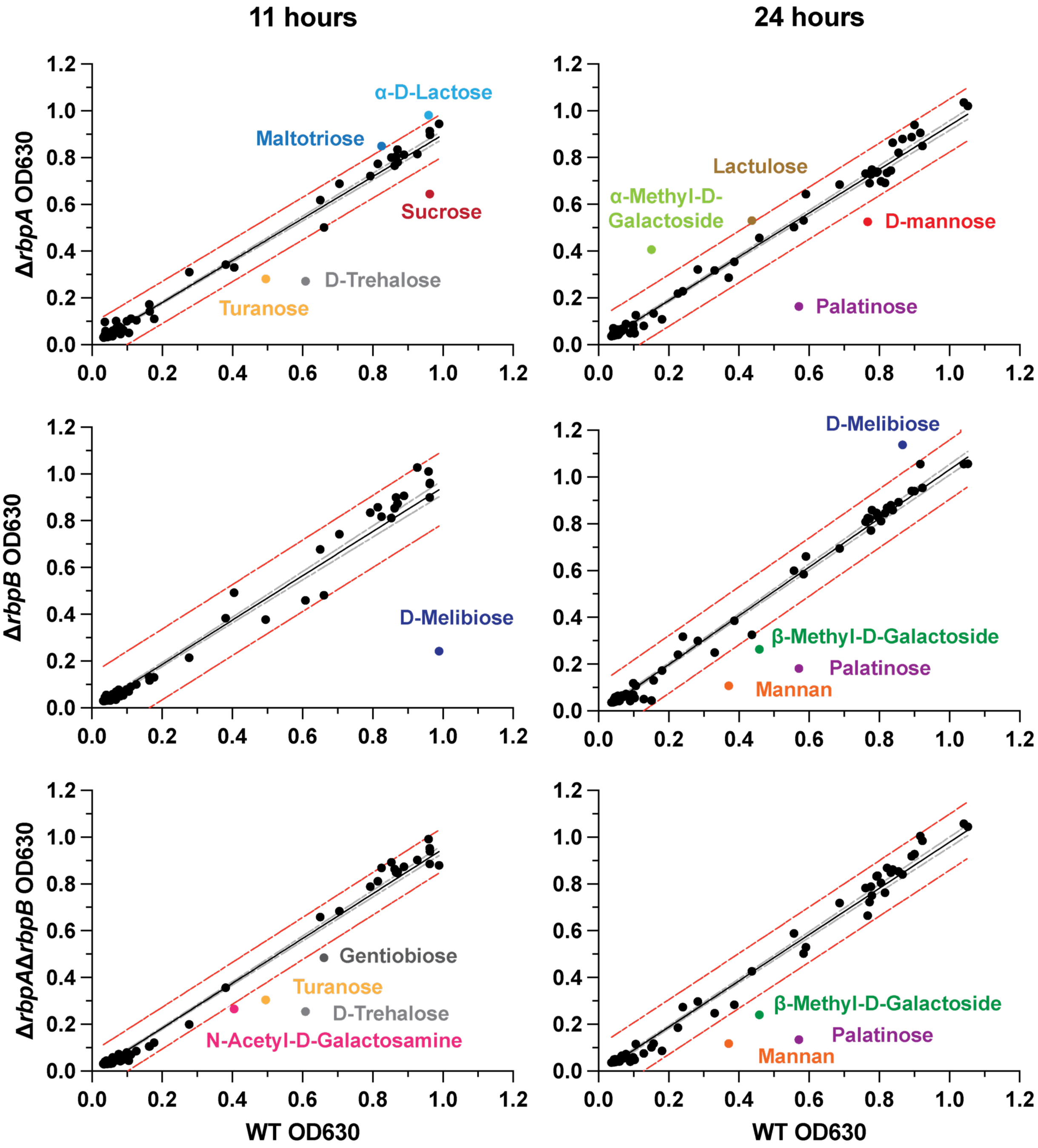


**Figure S2: Deletion of *rbpA* and *rbpB* perturbs growth on a variety of carbon sources.** Optical densities for WT, Δ*rbpA*, Δ*rbpB*, and Δ*rbpA*Δ*rbpB* at 11 hours of growth and 24 hours of growth on various carbon sources. WT values on the x-axis are the same for each mutant-WT comparison. The black line represents the linear regression and the grey line the regression 95% confidence interval. All carbohydrates outside the 99% prediction interval (red line) are highlighted. N=1.

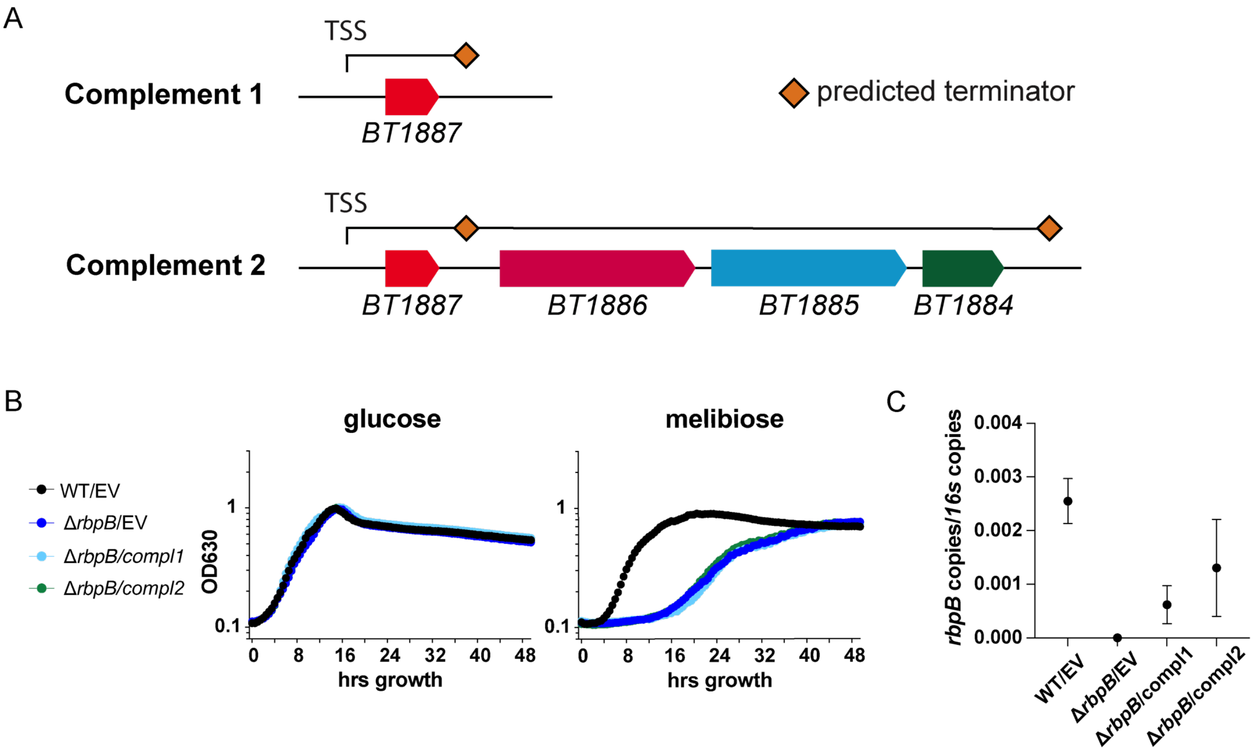


**Figure S3: Complementation with two *rbpB* constructs**. (A) Native genomic regions inserted into pNBU2 integration vectors for complementation of *rbpB (BT1887)* with transcription start site (TSS) and predicted terminators indicated. (B) Representative growth curves in minimal media from a single biological replicate (n=3) with optical densities ($OD_{630}$) recorded every 30 minutes. Empty Vector (EV) controls contain integrated pNBU2 without an insert. (C) RT-qPCR expression of *rbpB* complements during mid-log phase growth in MMG. The average and standard deviation of three biological replicates is plotted. *rbpB* mRNA levels from Δ*rbpB* and both complementation strains were significantly lower than WT (one-way ANOVA $F(3,8) = 12.85$) with Dunnett's multiple comparison's test, 8 DF, $p < 0.05$).

A

| | mid-log phase | | | stationary phase | | |
|---|---|---|---|---|---|---|
| | Δ*rbpA* | Δ*rbpB* | Δ*rbpA* Δ*rbpB* | Δ*rbpA* | Δ*rbpB* | Δ*rbpA* Δ*rbpB* |
| *BT1884* | NS | -1.54 | -3.63 | NS | NS | -1.59 |
| *BT1885* | NS | -2.08 | -2.72 | 1.80 | NS | NS |
| *BT1886* | NS | -2.20 | -2.93 | 1.75 | NS | 1.58 |
| *BT1887* | NS | -10.80 | -10.02 | NS | -7.83 | -8.68 |

| $log_2$ FC mut/WT |
|---|
| -12 |
| -8 |
| -4 |
| -2 |
| NS |
| 2 |

B

C

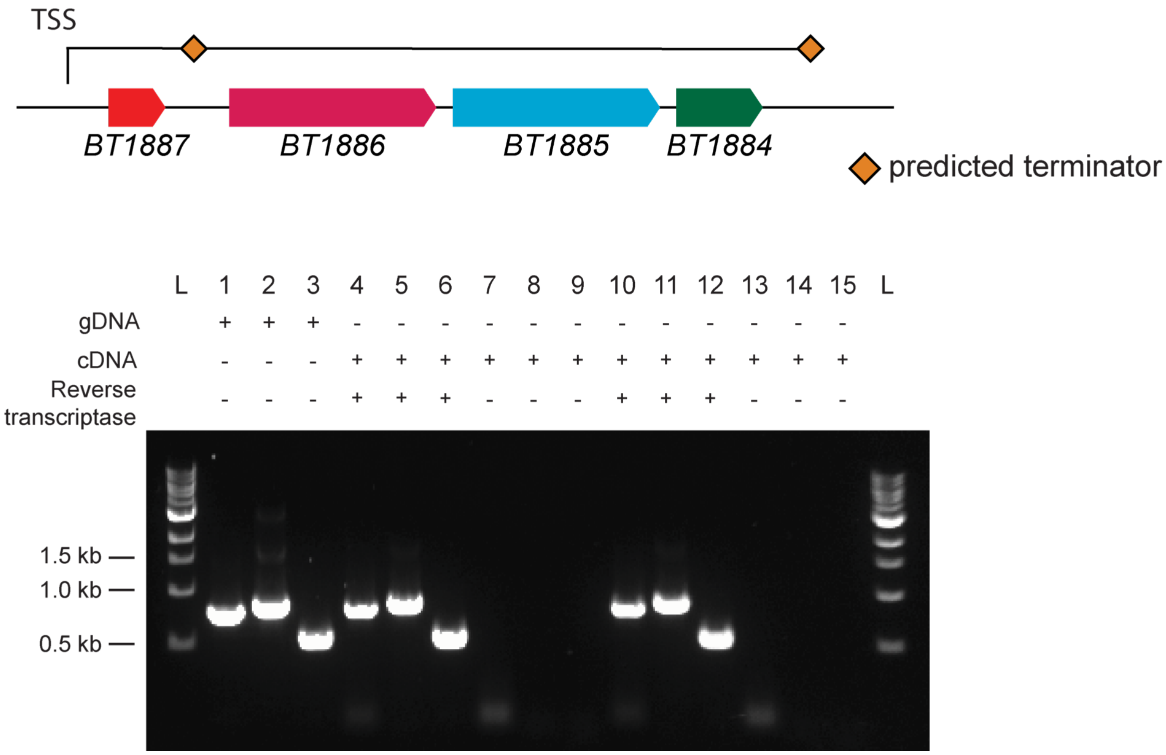


**Figure S4: *rbpB* is expressed in an operon with *BT1886*, *BT1885*, and *BT1884*.** (A) Expression values for genes *BT1884-BT1887* in TYG RNA-seq. Values shown had q-values <0.06. Comparisons that did not meet the q-value cutoff of <0.06 are labeled as Not Significant (NS). (B) Operon structure of *BT1887-BT1884*. Transcription Start Site (TSS) and predicted terminator positions indicated. (C) End-point RT-PCR of a representative example (n=3) of MMG (lanes 4-9) or MMM (lanes 10-15) cultures. Overlap regions amplified (lanes): *BT1884_BT1885* (1, 4, 7, 10, 13)), *BT1885_BT1886* (2, 5, 8, 11, 14), *BT1886_BT1887*(3, 6, 9, 12, 15). L=NEB 1Kb ladder.

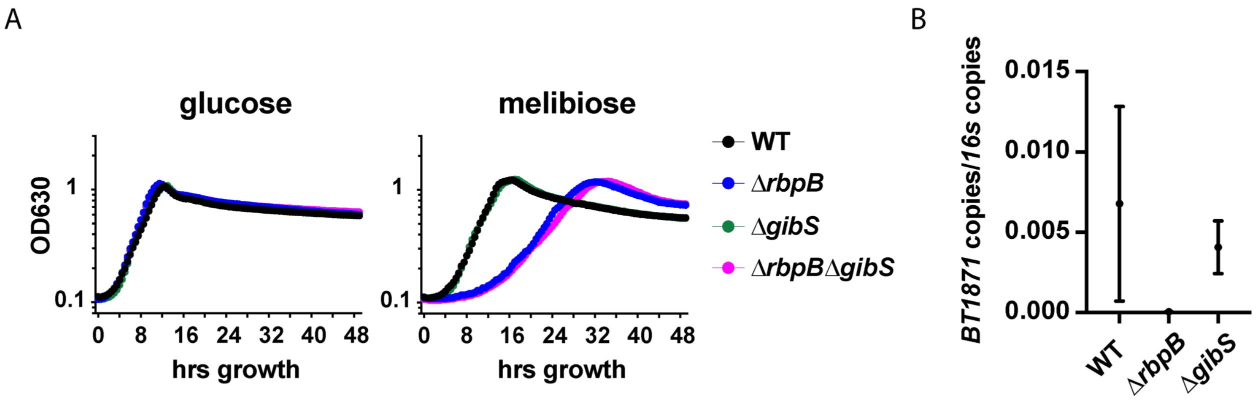


**Figure S5: Loss of *gibS* does not lead to melibiose growth defects**.
(A) Representative growth curves in minimal media from a single biological replicate (n=3) with optical densities ($OD_{630}$) recorded every 30 minutes. (B) RT-qPCR of *BT1871* mRNA from mid-log phase growth in MMG. The median and range of two biological replicates are plotted.

### Supplemental tables

**Table S1. Strains, plasmids, and oligonucleotides**

**Strains**

| Strain ID for this study | Bacterial strain | Genotype | Description | Source |
|---|---|---|---|---|
| S17-1 λ pir | *E. coli* (Migula) Castellani and Chalmers | *thi pro hdsR hdsM*+ *recA*, chromosomal insertion of RP4-2 (Tc::Mu Km::Tn7), $Amp^S$ | *E. coli* transformation strain used to conjugate plasmids into *B. thetaiotaomicron* VPI-5482. ATCC strain 47055. | (1) |
| AA0014 | *Bacteroides thetaiotaomicron* VPI-5482 | Δ*tdk* | Wildtype deletion background for *tdk* (*BT2275*)-based gene knockout method | (2) |
| AA0995 | *Bacteroides thetaiotaomicron* VPI-5482 | Δ*tdk*Δ*rbpA* | *rbpA (BT0784)* deletion strain made with pAA0991 in background strain AA0014 | This study |
| AA0671 | *Bacteroides thetaiotaomicron* VPI-5482 | Δ*tdk*Δ*rbpB* | *rbpB (BT1887)* deletion strain made with pAA1656 in background strain AA0014 | This study |
| AA0673 | *Bacteroides thetaiotaomicron* VPI-5482 | Δ*tdk*Δ*rbpA*Δ*rbpB* | *rbpA/rbpB* double knockout strain made with pAA0991 to delete *rbpA* in strain AA0014. *rbpA* deletion was then followed by *rbpB* deletion with pAA1656. | This study |
| AA1774 | *Bacteroides thetaiotaomicron* VPI-5482 | Δ*tdk*Δ*BT1871* | *BT1871* deletion strain made with pAA1738 in AA0014 | This study |
| AA1933 | *Bacteroides thetaiotaomicron* VPI-5482 | Δ*tdk*Δ*gibS* | *gibS* deletion strain made with pAA1909 in AA0014 | This study |
| AA1923 | *Bacteroides thetaiotaomicron* VPI-5482 | Δ*tdk*Δ*rbpB*Δ*gibS* | *gibS* deletion strain made with pAA1909 in AA0671 | This study |
| AA1963 | *Bacteroides thetaiotaomicron* VPI-5482 | Δ*tdk*_att1::pNBU2 | pAA0088 integrated in attachment site 1 of AA0014 | This study |
| AA1967 | *Bacteroides thetaiotaomicron* VPI-5482 | Δ*tdk*Δ*rbpB*_att1::pNBU2 | pAA0088 integrated in attachment site 1 of AA0671 | This study |
| AA1965 | *Bacteroides thetaiotaomicron* VPI-5482 | Δ*tdk*Δ*BT1871*_att1::pNBU2 | pAA0088 integrated in attachment site 1 of AA1774 | This study |
| AA1959 | *Bacteroides thetaiotaomicron* VPI-5482 | Δ*tdk*Δ*rbpB*_att1::pNBU2_*rbpB* | pAA1931 integrated in attachment site 1 of AA0671 | This study |
| AA1961 | *Bacteroides thetaiotaomicron* VPI-5482 | Δ*tdk*Δ*rbpB*_pNBU2_*rbpB-BT1884* | pAA1944 integrated in AA0671. Not in attachment sites 1 or 2. Undefined integration site. | This study |

| | | | | |
|---|---|---|---|---|
| AA1971 | *Bacteroides thetaiotaomicron* VPI-5482 | Δ*tdk*Δ*BT1871*_pNBU2_*BT1872-BT1871* | pAA1939 integrated in AA1774. Not in attachment sites 1 or 2. Undefined integration site. | This study |
| AA1969 | *Bacteroides thetaiotaomicron* VPI-5482 | Δ*tdk*Δ*BT1871*_att1::pNBU2_*BT1871* | pAA1943 integrated in attachment site 1 of AA1774 | This study |

**Plasmids**

| Plasmid ID for this study | Plasmid name extended | Description | Source |
|---|---|---|---|
| pET28b-*rpsA* | pET28b-*rpsA* | parent vector for pET28b-*rbpB* | (3) |
| pAA1836 | pET-*rbpB* | T7p-lacO-*rbpB*-His6 | This study |
| pAA0086 | pExchange_*bla_tdk_ermG* | parent vector for tdk suicide constructs | (2) |
| pAA0991 | pExchange_*rbpA* | *rbpA* (*BT0784*) suicide knockout vector | This study |
| pAA1656 | pExchange_*rbpB* | *rbpB* (*BT1887*) suicide knockout vector | This study |
| pAA1738 | pExchange_*BT1871* | *BT1871* suicide knockout vector | This study |
| pAA1824 | pLGB13_*bla_ermG* | parent vector for *gibS* suicide vector | (4) |
| pAA1909 | pLGB13_*gibS*_KO | *gibS* suicide knockout vector | This study |
| pAA0088 | pNBU2_*bla_ermG* | parent vector for complements and empty vector for complement controls | (5) |
| pAA1931 | pNBU2_*rbpB* | *rbpB* complement vector. Includes native 5' and 3' regions including 607 bp upstream of *rbpB* start codon and 521 bp downstream of r*bpB* stop codon. | This study |
| pAA1944 | pNBU2_*rbpB-BT1884* | *rbpB* through *BT1884* operon complement vector. Includes native 5' and 3' regions to the *rbpB-BT1884* operon including 600 bp upstream of *rbpB* start codon and 276 bp downstream of the *BT1884* stop codon. | This study |
| pAA1939 | pNBU2_*BT1872-BT1871* | *BT1872 through BT1871* full operon complement vector. Includes native 5' and 3' regions including 600 bp upstream of *BT1872* start codon and 879 bp downstream of the *BT1871* stop codon. | This study |
| pAA1943 | pNBU2_*BT1871* | *BT1871* only complement vector. Includes native 5' and 3' regions including 231 bp upstream of the *BT1871* start codon with a BT1872 ORF deletion. *BT1872* start and stop codons retained. Also includes 879 bp downstream of the *BT1871* stop codon. | This study |
| pAA1911 | pNBU2_*BT1872* | *BT1872* only complement vector. Includes 600 bp upstream of the *BT1872* start codon through the *BT1872* stop codon. | This study |

**Oligonucleotides**

| Oligo name for this study | Description (all are DNA oligos unless stated otherwise) | Sequence (5´-3´) | Source |
|---|---|---|---|
| OSA831 | Used to build pET-*rbpB*. Amplification of *rbpB* ORF for cloning into pET-28. | CTTTAAGAAGGAGATATACCATGAACATTTAC ATTTCAGGTTTAAGTTATGGCACAA | This study |

| | | | |
|---|---|---|---|
| OSA832 | Used to build pET-*rbpB*. Amplification of *rbpB* ORF for cloning into pET-28. | GATGATGGCTGCTGCCGCGCGGCACCAGAT ATCTTCTTGAATTGTTATAACCACCTCCACG | This study |
| OSA827 | Used to build pET-*rbpB*. Amplification of pET28 for assembly with *rbpB* ORF. | GGTATATCTCCTTCTTAAAGTTAAACAAAATT ATTTCTAGAGGGGAATT | This study |
| OSA828 | Used to build pET-*rbpB*. Amplification of pET28 for assembly with *rbpB* ORF. | CGCGGCAGCAGCCATCATC | This study |
| O-PP1 | ssDNA oligo from IDT, serves as template for dsDNA generation for PP1 and PP7 pentaprobe generation (complementary strand) | CGGAATTCTACGAATTTTTCTTTTGTTTATTTC CTTTCGCTTTGCTTCTCTTCCCTTCGTTCTGT TCCGTTTTACCTTGTCTTGCCTTATCTTACTT TA | (6) |
| O-PP2 | ssDNA oligo from IDT, serves as template for dsDNA generation for PP2 and PP8 pentaprobe generation (complementary strand) | TATCTTACTTTAGTTTCATTTAATTGTGTTGTA CTCTCCTCTGCGTTCACTTAGCTTAACTTGGT TTGGCTTGATTTGACTTCAGTTGCGCTCTATT CTA | (6) |
| O-PP3 | ssDNA oligo from IDT, serves as template for dsDNA generation for PP3 and PP9 pentaprobe generation (complementary strand) | CGCTCTATTCTACTGTCCTGTGCATTCAATC GTTGAGTTCGATCTAGTCTCGTCTAACCCTC CCCTGCTCCGCTGGTCTGGCCTCGCCTATC CTACCCAT | (6) |
| O-PP4 | ssDNA oligo from IDT, serves as template for dsDNA generation for PP4 and PP10 pentaprobe generation (complementary strand) | TATCCTACCCATTGGGCTCATCTGATCCATC CGGTCCCGTCCACTCGGCTATGTTATGCTGT ATTGCAGTCGTGTCGCGTCGAGCTGCCCTAA TCCCACC | (6) |
| O-PP5 | ssDNA oligo from IDT, serves as template for dsDNA generation for PP5 and PP11 pentaprobe generation (complementary strand) | CTAATCCCACCTAGCGTATCGGGTCATGTAG TGCTACGTTACGGCCCCCGCCCGGCATCAT ATTATATCACCCCAGTGTAATGTGGTGTGAG GTTGGAG | (6) |
| O-PP6 | ssDNA oligo from IDT, serves as template for dsDNA generation for PP6 and PP12 pentaprobe generation (complementary strand) | GTGAGGTTGGAGTCCGACCTGGAATCTCAG CCTGACGTGCCATGCGGTGCGATGTCACGC CGCGCCACGGTATAGTATGGTACGGGATCC CG | (6) |
| OSA857 | T7 forward primer for amplification of O-PP1, produces PP1 dsDNA reverse transcription template | TAATACGACTCACTATAGGGCGGAATTCTAC GAATTTTTCTTTTGTTTATTTCCTTTC | (6) |
| OSA858 | Reverse primer for amplification of O-PP1, produces PP1 dsDNA reverse transcription template | TAAAGTAAGATAAGGCAAGACAAGGTAAAAC GG | (6) |
| OSA859 | T7 forward primer for amplification of O-PP1, produces PP7 dsDNA reverse transcription template | TAATACGACTCACTATAGGGTAAAGTAAGAT AAGGCAAGACAAGGTAAAACGG | (6) |
| OSA860 | Reverse primer for amplification of O-PP1, produces PP7 dsDNA reverse transcription template | TAATACGACTCACTATAGGGCGGAATTCTAC GAATTTTTCTTTTGTTTATTTCCTTTC | (6) |
| OSA861 | T7 forward primer for amplification of O-PP2, produces PP2 dsDNA reverse transcription template | TAAAGTAAGATAAGGCAAGACAAGGTAAAAC GG | (6) |
| OSA862 | Reverse primer for amplification of O-PP2, produces PP2 dsDNA reverse transcription template | TAATACGACTCACTATAGGGTAAAGTAAGAT AAGGCAAGACAAGGTAAAACGG | (6) |
| OSA857 | T7 forward primer for amplification of O-PP2, produces PP8 dsDNA reverse transcription template | CGGAATTCTACGAATTTTTCTTTTGTTTATTTC CTTTC | (6) |
| OSA858 | Reverse primer for amplification of O-PP2, produces PP8 dsDNA reverse transcription template | TAATACGACTCACTATAGGGTATCTTACTTTA GTTTCATTTAATTGTGTTGTACTCTCCT | (6) |

| | | | |
|---|---|---|---|
| OSA859 | T7 forward primer for amplification of O-PP3, produces PP3 dsDNA reverse transcription template | TAGAATAGAGCGCAACTGAAGTCAAATCA | (6) |
| OSA860 | Reverse primer for amplification of O-PP3, produces PP3 dsDNA reverse transcription template | TAATACGACTCACTATAGGGTAGAATAGAGCGCAACTGAAGTCAAATCA | (6) |
| OSA861 | T7 forward primer for amplification of O-PP3, produces PP9 dsDNA reverse transcription template | TATCTTACTTTAGTTTCATTTAATTGTGTTGTACTCTCCT | (6) |
| OSA862 | Reverse primer for amplification of O-PP3, produces PP9 dsDNA reverse transcription template | TAATACGACTCACTATAGGGCGCTCTATTCTACTGTCCTGTGCA | (6) |
| OSA863 | T7 forward primer for amplification of O-PP4, produces PP8 dsDNA reverse transcription template | ATGGGTAGGATAGGCGAGGC | (6) |
| OSA864 | Reverse primer for amplification of O-PP4, produces PP8 dsDNA reverse transcription template | TAATACGACTCACTATAGGGATGGGTAGGATAGGCGAGGC | (6) |
| OSA865 | T7 forward primer for amplification of O-PP4, produces PP10 dsDNA reverse transcription template | CGCTCTATTCTACTGTCCTGTGCA | (6) |
| OSA866 | Reverse primer for amplificationo of O-PP4, produces PP10 dsDNA reverse transcription template | TAATACGACTCACTATAGGGTATCCTACCCATTGGGCTCATCTGAT | (6) |
| OSA867 | T7 forward primer for amplification of O-PP5, produces PP5 dsDNA reverse transcription template | GGTGGGATTAGGGCAGCTCG | (6) |
| OSA868 | Reverse primer for amplification of O-PP5, produces PP5 dsDNA reverse transcription template | TAATACGACTCACTATAGGGGGTGGGATTAGGGCAGCTCG | (6) |
| OSA869 | T7 forward primer for amplification of O-PP5, produces PP11 dsDNA reverse transcription template | TATCCTACCCATTGGGCTCATCTGAT | (6) |
| OSA870 | Reverse primer for amplification of O-PP5, produces PP11 dsDNA reverse transcription template | TAATACGACTCACTATAGGGCTAATCCCACCTAGCGTATCGGG | (6) |
| OSA871 | T7 forward primer for amplification of O-PP6, produces PP6 dsDNA reverse transcription template | CTCCAACCTCACACCACATTACAC | (6) |
| OSA872 | Reverse primer for amplification of O-PP6, produces PP6 dsDNA reverse transcription template | TAATACGACTCACTATAGGGCTCCAACCTCACACCACATTACAC | (6) |
| OSA873 | T7 forward primer for amplification of O-PP6, produces PP12 dsDNA reverse transcription template | CTAATCCCACCTAGCGTATCGGG | (6) |
| OSA874 | Reverse primer for amplification of O-PP6, produces PP12 dsDNA reverse transcription template | TAATACGACTCACTATAGGGGTGAGGTTGGAGTCCGACCT | (6) |
| Motif1 | Motif 1 rRNA used for EMSAs with RbpB | GUAGGAUAGUAGGAUAGUAGGAUA | This study |
| Motif9 | Motif 9 rRNA used for EMSAs with RbpB | UCCUGUGCUCCUGUGCUCCUGUGC | This study |
| A.C02 | BT1883_BT1884 RT-PCR forward primer | GGTGCACCTTCCCACCATAACGG | This study |
| A.C03 | BT1883_BT1884 RT-PCR reverse primer | CGAGGAACCCGTTATATTGAGAGGGAGAG | This study |

| | | | |
|---|---|---|---|
| A.C04 | BT1884_BT1885 RT-PCR forward primer | CATTCTCGTCCACATAAGCAATCATATCA | This study |
| A.C05 | BT1884_BT1885 RT-PCR reverse primer | GCTACGATGCCTGATACGATTATTGCTCTG | This study |
| A.C06 | BT1885_BT1886 RT-PCR forward primer | CAGAGCAATAATCGTATCAGGCATCGTAGC | This study |
| A.C07 | BT1885_BT1886 RT-PCR reverse primer | GCATTATTAAAGCGCTGTATAATGCGGCC | This study |
| A.C08 | BT1886_BT1887 RT-PCR forward primer | CCCCCTTCGTTTGCTACTGCTGAC | This study |
| A.C60 | BT1886_BT1887 RT-PCR reverse primer | CGGTGGTAACCGTGGAGGTGG | This study |
| A.C59 | BT1887_BT1888 RT-PCR forward primer | CCACCTCCACGGTTACCACCG | This study |
| A.C09 | BT1887_BT1888 RT-PCR reverse primer | GCTTTGCCCACAATTCTTCCCGTGTG | This study |
| A.D11 | 16s qPCR forward primer | GGTAGTCCACACAGTAAACGATGAA | (7) |
| A.D12 | 16s qPCR reverse primer | CCCGTCAATTCCTTTGAGTTTC | (7) |
| A.D13 | BT1887 qPCR forward primer | TAGAGAAACCGGCAGATCCA | This study |
| A.D14 | BT1887 qPCR forward primer | GGTTACCACCGTATGACG | This study |
| NBU2_att1_F | pNBU2 integrase attachment site 1 check forward primer | CCTTTGCACCGCTTTCAACG | (5) |
| NBU2_att1_R | pNBU2 integrase attachment site 1 check reverse primer | TCAACTAAACATGAGATACTAGC | (5) |
| NBU2_att2_F | pNBU2 integrase attachment site 2 check forward primer | TATCCTATTCTTTAGAGCGCAC | (5) |
| NBU2_att2_R | pNBU2 integrase attachment site 2 check reverse primer | GGTGTACCTGGCATTGAAGG | (5) |
| ZC_rbpA_LF_BamHI | Used to build pAA0991. Forward primer for amplification of upstream *BT0784* flank (left flank) with ZC_BT0784_LF_SOE from the genome. Contains BamHI site for assembly into pAA0086. | GTCCGCGGATCCCCTTTAGAGATACGAACGTCGGC | This study |
| ZC_rbpA_LF_SOE | Used to build pAA0991. Reverse primer for amplification of upstream *BT0784* flank with ZC_BT0784_LF_BamHI from the genome. Contains overlapping region for SOE with *BT0784* downstream region (right flank). | CCTTCTTTTTATATACTGATTGTTATTCAGTCAATTACATAAAATACAAATATAAAAAATTAATAATTTCAATAACAAAG | This study |
| ZC_rbpA_RF_SOE | Used to build pAA0991. Forward primer for amplification of downstream *BT0784* flank (right flank) with ZC_BT0784_RF_NotI. Contains overlapping region for SOE with *BT0784* left flank. | GAAATTATTAATTTTTTATATTTGTATTTTATGTAATTGACTGAATAACAATCAGTATATAAAAAGAAGGTGTG | This study |

| | | | |
|---|---|---|---|
| ZC_rbpA_RF_NotI | Used to build pAA0991. Reverse primer for amplification of downstream *BT0784* flank (right flank) with ZC_BT0784_RF_SOE from the genome. Contains NotI site for assembly into pAA0086. | ATGTCGCGGCCGCGTAATCTTAGGCGCGAATGTC | This study |
| ZC_rbpB_LF_BamHI | Used to build pAA1656. Forward primer for amplification of upstream *BT1887* flank (left flank) with ZC_BT1887_LF_SOE from the genome. Contains BamHI site for assembly into pAA0086. | GGCAGGGATCCGCCATTTGTCACAGATTATTTTGGCAG | This study |
| ZC_rbpB_LF_SOE | Used to build pAA1656. Reverse primer for amplification of upstream *BT1887* flank with ZC_BT1887_LF_BamHI from the genome. Contains overlapping region for SOE with *BT1887* downstream region (right flank). | CAAATTAGTATTTATTTTTAATAAAATTTTAGATGTAATATCAGAATAAATAAAAGCAGAAGATAATGTCTTCTG | This study |
| ZC_rbpB_RF_SOE | Used to build pAA1656. Forward primer for amplification of downstream *BT1887* flank (right flank) with ZC_BT1887_RF_NotI from the genome. Contains overlapping region for SOE with *BT1887* left flank. | CATTATCTTCTGCTTTTATTTATTCTGATATTACATCTAAAATTTTATTAAAAATAAATACTAATTTGGAGTGAGGAGG | This study |
| ZC_rbpB_RF_NotI | Used to build pAA1656. Reverse primer for amplification of downstream *BT1887* flank (right flank) with ZC_BT1887_RF_SOE from the genome. Contains NotI site for assembly into pAA0086. | GTCTGCGCGGCCGCGCTTTTCCATCGATGGTCTGC | This study |
| OXM262 | Used to build pAA1943. Forward primer used with OXM263 for amplification of the pAA1943 full insert after OXM263-OXM268 PCR fragment SOE assembly with OXM267. Contains NotI site for assembly into pAA0088. | AATTATGCGGCCGCTGTGGCTTTTTCTTTCTGAACCGTCTATTG | This study |
| OXM263 | Used to build pAA1943. Reverse primer for amplifying *BT1871* from the genome with OXM268. OXM263-OXM268 PCR fragment was then assembled with gBlock OXM267 using SOE. OXM268 also used with OXM263 to amplify the full SOE insert for assembly into pAA0088. Contains PstI site for assembly into pAA0088. | AATTATCTGCAGATGCATAAATTTGCAAGCACCAACAAAAG | This study |
| OXM268 | Used to build pAA1943. Forward primer for amplifying *BT1871* from the genome with OXM263. | ATGAAAAAACTTACATTTTTATTATTATGTGTCTTGTGTAC | This study |

| | | | |
|---|---|---|---|
| OXM349 | Used to build pAA1909. Forward primer for amplifying left flank to *gibS* TSS with OXM350. Contains BamHI site. | AATTATGGATCCGGAAGGCGACAGAATGTATTTTAAGAG | This study |
| OXM350 | Used to build pAA1909. Reverse primer for amplifying left flank to *gibS* TSS with OXM349. | ACTACAAAACGCAACAAAAGGACATAAATATAACAGATTAAAAAGAGGCTTTGTTCAAC | This study |
| OXM351 | Used to build pAA1909. Forward primer for amplifying right flank to *gibS* with OXM352. | GTTGAACAAAGCCTCTTTTTAATCTGTTATATTTATGTCCTTTTGTTGCGTTTTGTAGT | This study |
| OXM352 | Used to build pAA1909. Reverse primer for amplifying right flank to *gibS* with OXM351. Contains SalI site. | AATTATGTCGACGGTATTATTTTATTCAAAATCATCAATTGTGTAATTGATG | This study |
| OXM365 | Used to build pAA1939 and pAA1911. Forward primer for amplification of BT1872 and BT1871 operon for cloning into pAA0088 for pAA1939. Forward primer for amplification of BT1872 for cloning into pAA0088 for pAA1911. Contains XbaI site. | AATTATTCTAGATGGCGAAAGATCTGAAAACGACAGAAGGTG | This study |
| OXM366 | Used to build pAA1939. Reverse primer for amplification of BT1872 and BT1871 operon for cloning into pAA0088. Contains BamHI site. | AATTATGGATCCATGCATAAATTTGCAAGCACCAACAAAAGACAACTAAC | This study |
| OXM367 | Used to build pAA1911. Reverse primer for amplification of BT1872 for cloning into pAA0088 with BamHI. Contains BamHI site. | AATTATGGATCCTTATTCCGCAATAAAGCGCTTTGTCTGTACATCAC | This study |
| OXM379 | Used to build pAA1931 and pAA1944. Forward primer used with OXM380 to amplifiy *BT1887* (*rbpB*) through *BT1884* 3' UTR from the genome to make pAA1944. Also used as a forward primer with OXM381 to amplify *BT1887* to make pAA1931. Contains BamHI site for assembly into pAA0088. | AATTATGGATCCTCGTCGGGATGGATGCGCGATTC | This study |
| OXM380 | Used to build pAA1944. Reverse primer used with OXM379 to amplify *BT1887* through *BT1884* 3' UTR from the genome. Contains PstI site for assembly into pAA0088. | AATTATCTGCAGGTTTTCTGCGTTTAAAAGTTAATAATATTTCAGACTCAGCGGGCA | This study |
| OXM381 | Used to build pAA1931. Reverse primer used with OXM378 to amplify *BT1887* from the genome. Contains PstI site for assembly into pAA0088. | AATTATCTGCAGTTGGGATAAAGCAACCCCAGACCTG | This study |

| | | | |
|---|---|---|---|
| BT1871_KO | IDT gBlock used to build pAA1738. Fragment contains flanks to *BT1871* ORF. Contains digest sites for BamHI (5' digest) and XbaI (3' digest). | GACGGGGATCCATCTGGAAATCCCTGACGC<br>TCAGCAGGACTTATTGAAAGCATTGGTGAAA<br>ACAGGTAAGCCTGTGGTACTGCTCCTGTTCA<br>CCGGTCGTCCTTTGATCTTGAACTGGGAAAG<br>CGAACACATTCCTTCTATTCTGAATGTATGGT<br>TCGGAGGTAGTGAGACAGGGGATGCCGTTG<br>CCGATGTTCTGTTTGGCAAAGCCGTCCCGTG<br>CGGTAAGCTGACTACTACCTTCCCTCGTTCG<br>GTAGGGCAACTTCCTTTGTTCTACAATCATTT<br>GAACACTGGACGTCCGGACCCGGACAACCG<br>TGTTTTCAACCGGTATGCCAGCAACTATCTG<br>GATGAGAGTAACGAACCTCTTTATCCTTTCG<br>GATACGGTTTGAGTTATACCGACTTTGTATAT<br>GGCGACCTGCAGCTCAGTTCGGAAACCTTG<br>CCGAAAAATGGAAACCTGACAGCTTCCGTTA<br>CCGTCACCAACAAAGGGAACCATGACGGCT<br>ATGAAACGGTACAAATCTATTTGCGTGATATC<br>TATGCGGAAGTAGCGCGTCCGGTGAAAGAG<br>CTGAAAGGCTTCGATCGTATCTTCCTGAAAA<br>AAGGAGAAAGCCGTGAAGTGAAGTTTGTACT<br>TACGGAAGATGACCTGAAATTTTATAATTCCG<br>GGCTGCAATATATCTACGAACCGGGAGAGTT<br>TGACGTGATGATAGGTACGAACAGTCGTGAT<br>GTACAGACAAAGCGCTTTATTGCGGAATAAT<br>TATTTTGAAATCTAATATATTAGTTACAACCAT<br>GTAATTATTTTAGAGTTGAAACTATGTTTTTTA<br>TCGTCACCCGTCACTTTTTAGAATAAAGTATT<br>AATATATAGACCAATAGCGGTGACGGCAAAT<br>GCTGATGGCTAAGTGACGGCAAGAGTTTTAC<br>GCTTTTTGCCGTCACTTTTTGTGTTGTAAACT<br>GATTCTTGGTCTAGCATATTATTCTCCTAGAG<br>ATAATCAGGCTCCCCACGTAATTTGAATGCG<br>TTTTTGCTCTCATGCCCTAACAGATTGTCTGT<br>ATGCCTTTATTCATCGTAGTTTCAGGTATGGC<br>AGCAATTGTGGTAGCAGGTATGAGAGCAAGT<br>GTTTGCTCTCACTCTTCCATGTATCATTATCT<br>GTCATTTGTTTGATGTTGTCATTTGATTTACA<br>GTGTGTTAATGTCTGAGGTGTCAATTCTTAAC<br>GATTGATATTTCAGACATTAATAATTGATAAA<br>TCATCCTTTAATAGTTGATAACTCAGCTGTTT<br>TTGAGTGATAAGTGATTTATTGTTTCTTGGAA<br>GAGCAGGTTAATCGAACAGTTGGTGCGGTC<br>AAGTTAATTGAATCATATAAGTAAGGGATCAG<br>TTTCCTGCCTTGTTTCATTCATTCCATCGGCT<br>GGAACCACTGTTTTCAAGCGTCCGGAACTTT<br>GGTTTCACGGCTTTGAAACTAAAGTTTCAAG<br>CCGATGAAACTTTAGTTCCAAGCAGGAGAAA<br>CAAAAGTTTCAAGCGGGAGAAACTAAAAAGC<br>TCTGTCTAGACATCA | This study |
| OXM267 | IDT gBlock used to build pAA1943. Fragment contains the native promoter region for *BT1871* without the *BT1872* ORF. Contains NotI digest site for assembly into pAA0088. | GCGGCCGCTGTGGCTTTTTCTTTCTGAACCG<br>TCTATTGTATATAGATTCAGTTGACAGAAATA<br>AATAATCTATTTTGCGGGTATATGTCTCATTA<br>TTTAGAAGGAAGTGTTAACTTTGGCGAAATA<br>GATTCTGTTGACCGATGTTGCCGTTATTGTAT<br>AGAAGGTAAAGTCAGAAAAGTAAATAAATAAT<br>TTGAAGAGCTTATGTAATTATTTTGAAATCTA<br>ATATATTAGTTACAACCATGAAAAAACTTACA<br>TTTTTATTATTATGTGTCTTGTGTACGCTTTC<br>GTTGCAAGCTCAGAAGCAATTTACACTAGCC<br>TCACCTGATGGAAATCTGAAAACGAC | This study |

**Table S2: Publicly available RNA-seq datasets used to identify RBP expression *in vitro* and *in vivo***

| Strain | *rbp* genes | Media, carbon source, or diet | Sample ID | Citation |
|---|---|---|---|---|
| *B. thetaiotaomicron* VPI-5482 | *BT0784 (rbpA)*<br>*BT1887 (rbpB)*<br>*BT3840 (rbpC)* | *In vitro*<br>minimal media glucose | SRR5973480<br>SRR5973481 | (1) |
| *B. thetaiotaomicron* VPI-5482 | | *In vitro*<br>TYG | SRR14277227<br>SRR14277226 | Unpublished |
| *B. thetaiotaomicron* VPI-5482 | | *In vivo*<br>monocolonized, gnotobiotic mouse cecum, standard mouse chow (5K67 LabDiet, Purina) | ERS2154079<br>ERS2154080<br>ERS2154081 | (2) |
| *B. thetaiotaomicron* VPI-5482 | | *In vivo*<br>synthetic community gnotbiotic mouse cecum, fiber-rich chow (LabDiet 5010, Purina) | SRR4838357<br>SRR4838358<br>SRR4838359 | (3) |
| *B. caccae* ATCC 43185 | *BACCAC00656 (rbpA)*<br>*BACCAC03695 (rbpB)*<br>*BACCAC03375 (rbpC)* | *In vitro*<br>minimal media glucose | SRR4876883<br>SRR4876890 | (3) |
| *B. cellulosilyticus* WH2 | *BWH2RS0117595 (rbpA)*<br>*BWH2RS0107135 (rbpB)*<br>*BWH2RS0123290 (rbpC)*<br>*BWH2RS0115465 (rbpF)* | *In vitro*<br>TYG | ERR922690<br>ERR922691<br>ERR922692<br>ERR922693 | (4) |
| *B. dorei* ATCC 8492 | *BACDOR02742 (rbpA)*<br>*BACDOR01223 (rbpA')*<br>*BACDOR00009 (rbpC)*<br>*BACDOR02324 (rbpD)* | *In vitro*<br>GIFU glucose | SRR10196317<br>SRR10196295<br>SRR10196273 | (5) |
| *B. fragilis* NCTC 9343 | *BFnctc2225 (rbpA)*<br>*BFnctc2361 (rbpB)*<br>*BFnctc3897 (rbpC)*<br>*BFnctc2734 (rbpD)* | *In vitro*<br>minimal media glucose | SRR2584375<br>SRR2585000<br>SRR2602448 | (6) |
| *B. massiliensis* DSM 17679 | *HMPREF1534_RS07415 (rbpC)*<br>*HMPREF1534_RS05480 (rbpD)* | *In vitro*<br>minimal media glucose | SRR2583239<br>SRR2583240<br>SRR2583566 | (6) |
| *B. uniformis* ATCC 8492 | *BACUNI03361 (rbpA)*<br>*BACUNI01393 (rbpB)*<br>*BACUNI03010 (rbpC)* | *In vitro*<br>minimal media glucose | SRR7216289<br>SRR7216288 | (7) |
| *B. vulgatus* ATCC 8482 | *BVU_3246 (rbpA)*<br>*BVU_0543 (rbpA')*<br>*BVU_0446 (rbpC)*<br>*BVU_1413 (rbpD)* | *In vitro*<br>minimal media glucose | SRR7216285<br>SRR7216284 | (7) |
| *B. xylanisolvens* XB1A | *Bxyl0536 (rbpA)*<br>*Bxyl4341 (rbpC)* | *In vitro*<br>Bx-medium | SRR2827524<br>SRR2827525<br>SRR2827526 | (8) |

1. Costliow ZA, Degnan PH. 2017. Thiamine acquisition strategies impact metabolism and competition in the gut microbe *Bacteroides thetaiotaomicron*. mSystems 2:e00116-17.

2. Schofield WB, Zimmermann-Kogadeeva M, Zimmermann M, Barry NA, Goodman AL. 2018. The stringent response determines the ability of a commensal bacterium to survive starvation and to persist in the gut. Cell Host Microbe 24:120–132.

3. Desai MS, Seekatz AM, Koropatkin NM, Kamada N, Hickey CA, Wolter M, Pudlo NA, Kitamoto S, Terrapon N, Muller A, Young VB, Henrissat B, Wilmes P, Stappenbeck TS, Núñez G, Martens EC. 2016. A dietary fiber-deprived gut microbiota degrades the colonic mucus barrier and enhances pathogen susceptibility. Cell 167:1339–1353.

4. Wu M, McNulty NP, Rodionov DA, Khoroshkin MS, Griffin NW, Cheng J, Latreille P, Kerstetter RA, Terrapon N, Henrissat B, Osterman AL, Gordon JI. 2015. Genetic determinants of *in vivo* fitness and diet responsiveness in multiple human gut *Bacteroides*. Science 350:aac5992.

5. Huang Y, Sheth RU, Kaufman A, Wang HH. 2019. Scalable and cost-effective ribonuclease-based rRNA depletion for transcriptomics. Nucleic Acids Res 48:e20.

6. Pudlo NA, Urs K, Kumar SS, German JB, Mills DA, Martens EC. 2015. Symbiotic human gut bacteria with variable metabolic priorities for host mucosal glycans. mBio 6:e01282-15.

7. Costliow ZA, Degnan PH, Vanderpool CK. 2019. Thiamine pyrophosphate riboswitches in *Bacteroides* species regulate transcription or translation of thiamine transport and biosynthesis genes. bioRxiv 867226.

8. Despres J, Forano E, Lepercq P, Comtet-Marre S, Jubelin G, Chambon C, Yeoman CJ, Miller MEB, Fields CJ, Martens E, Terrapon N, Henrissat B, White BA, Mosoni P. 2016. Xylan degradation by the human gut *Bacteroides xylanisolvens* XB1AT involves two distinct gene clusters that are linked at the transcriptional level. BMC Genomics 17:1–14.
